# The yeast phosphofructokinase β-subunit has ATP-dependent RNA unwinding activity and modulates cell cycle progression

**DOI:** 10.1101/2025.02.01.636022

**Authors:** Waleed S. Albihlal, Ana M. Matia-González, Tobias Schmidt, Wieland Mayer, Martin Bushell, Dierk Niessing, Matteo Barberis, Alexander Schmidt, Jürgen J. Heinisch, André P. Gerber

## Abstract

Phosphofructokinase (PFK) is a rate-limiting glycolytic enzyme that also possesses an unexplored RNA binding activity. Here, we show that the α- and β-subunits of yeast PFK, encoded by *PFK1* and *PFK2*, respectively, bind hundreds of mRNAs in cells, including one’s coding for proteins involved in regulation of mitotic cell cycle. Pkf2p directly binds in a Mg-ATP-dependent manner to short GA-, UC-, AU- and U-rich motifs overrepresented in its mRNA targets. Strikingly, Pfk2p displays directional 5’ - 3’ double-stranded RNA unwinding activity not seen with Pfk1p. Furthermore, Pfk2p dynamically associates with ribosomes and promotes translation of cell cycle genes. Consequently, *pfk2Δ,* but not *pfk1Δ,* mutant cells show severely delayed G1/S phase transition which is independent of the enzyme’s glycolytic activity. Our results uncovered a hidden function for the Pfk2 subunit as a translational activator of mitotic cell cycle gene transcripts possibly through energy-dependent RNA unwinding activity.

## Introduction

RNA binding proteins (RBPs) play vital roles in the post-transcriptional control of gene expression ^1,2^. The development of various proteomics approaches during the last decade has greatly expanded the repertoire of eukaryotic RNA-binding proteins (RBPs). Besides “conventional” RBPs, which possess one or more well-characterized or predicted RNA-binding domains (RBDs), hundreds of “unconventional” RBPs lacking defined RBDs but bearing other well-established cellular functions such as metabolic enzymes were identified ^1–3^. For instance, several early studies suggested that several glycolytic enzymes act as RBPs ^3,4^. More recently, crosslinking-assisted RNA interactome capture (RIC) studies revealed that most if not all glycolytic enzymes can interact with polyadenylated (poly(A)) RNA in the yeast *Saccharomyces cerevisiae* and in other eukaryotes, suggesting consequences for the bound RNAs, the interacting enzyme and/or for both ^3,5–8^.

Glycolysis is an evolutionarily conserved energy generating pathway breaking down glucose (glc) into pyruvate with net production of two molecules ATP and two molecules of NADH, which is an essential cofactor. Phosphofructokinase (PFK) is the ‘gate keeper’ of glycolysis and catalyses the irreversible conversion of fructose-6-phosphate (F6P) and ATP to fructose-1,6-bisphosphate (F1,6BP) and ADP. Importantly, PFK is highly allosterically regulated allowing for fine-tuning of the glycolytic flux and adjusting cellular energy demands: it is activated by AMP/ADP and fructose 2,6-biphosphate (F2,6BP), and inhibited by high levels of ATP, citrate, and other metabolites ^9,10^. Thereby, each subunit of eukaryotic Pfks consists of two homologous halves related to a potential bacterial ancestor: the N-terminal half containing substrate and ATP binding sites with catalytic functions, while the C-terminal half bears effector-binding sites for allosteric regulation ^11–13^.

In the yeast *Saccharomyces cerevisiae,* PFK is comprised of two paralogous subunits, termed α (Pfk1p) and β (Pfk2p) that can form a heterooctameric complex ^11,12,14–17^. While both yeast Pfk subunits cooperate in glycolysis, early mutational studies suggested additional and possibly distinct functions for both paralogs. For instance, deletion of *PFK2* leads to significantly slower cell growth, while deleting *PFK1* shows no noticeable effect on growth, despite the commonly observed lack of *in vitro* catalytic activity in the respective deletions or through mutation of the catalytic centre ^18–20^. Interestingly, a recent report suggested that the yeast Pfk subunits besides eight other glycolytic enzymes, assemble into cytoplasmic granules called glycolytic bodies (G-bodies) under hypoxic stress conditions; where they associated with numerous RNAs, including mRNAs coding for glycolytic enzymes ^7^. Moreover, photoactivatable ribonucleoside-enhanced crosslinking and immunoprecipitation (PAR-CLIP) followed by deep sequencing suggested that Pfk2p binds to numerous mRNAs that contain AU/U-rich elements at binding sites preferentially located in 3’UTRs, which further confirmed the RNA binding activity of Pfk2p though its functional relevance remained unclear ^7^.

Here we describe novel functions of the yeast Pfk2p in RNA metabolism under normal physiological growth conditions which complements its role in glycolysis. We found that Pfk proteins bind to hundreds of mRNAs coding for functionally related proteins including mitotic cell cycle regulators, in a glucose dependent manner *in vivo*. The bound mRNAs contain distinct short RNA triplet motifs preferentially located either in CDS or 3’UTRs; and Pfk2p binds to these motifs in an Mg-ATP dependent manner. Most intriguingly, Pfk2p exhibits RNA unwinding activity with high 5’ - 3’ polarity, and it promotes translation of cell cycle gene transcripts such as *cyclin 3* (*CLN3*). Consistently, *pfk2Δ*, but not *pfk1Δ*, mutant cells displayed delayed progression into S phase of the cell cycle which is independent of the enzyme’s catalytic activity. In conclusion, our results suggest a model in which Pfk2p acts as a molecular relay that controls cell cycle progression promoting the energy-dependent translation of cell cycle regulators.

## Results

### Pfk1p and Pfk2p bind RNA in vivo

To reconcile the RNA-binding activities of Pfk1p and Pfk2p seen in previous RIC-MS studies ^5,6^, we first performed RIC with yeast strains expressing either *PFK1* or *PFK2* with a C-terminal tandem-affinity purification (TAP)-tag under the control of their endogenous promoters. As expected, Pfk1:TAP and Pfk2:TAP were detected in RIC eluates but not in corresponding samples supplemented with a poly(A) competitor. No signals were detected for Act1p, a non-RBP control (Figure 1A). Likewise experiments with wild-type cells confirmed interaction of both endogenous (untagged) Pfk1 and Pfk2 proteins, using an antibody that detects both Pfk subunits. RIC performed with *pfk1*Δ and *pfk2*Δ strains further revealed that each subunit can interact with poly(A)-RNA in the absence of the other, indicating that formation of an heterooctameric Pfk protein complex is not required for interaction with poly(A) RNAs (Figure 1A).

**Figure 1.**
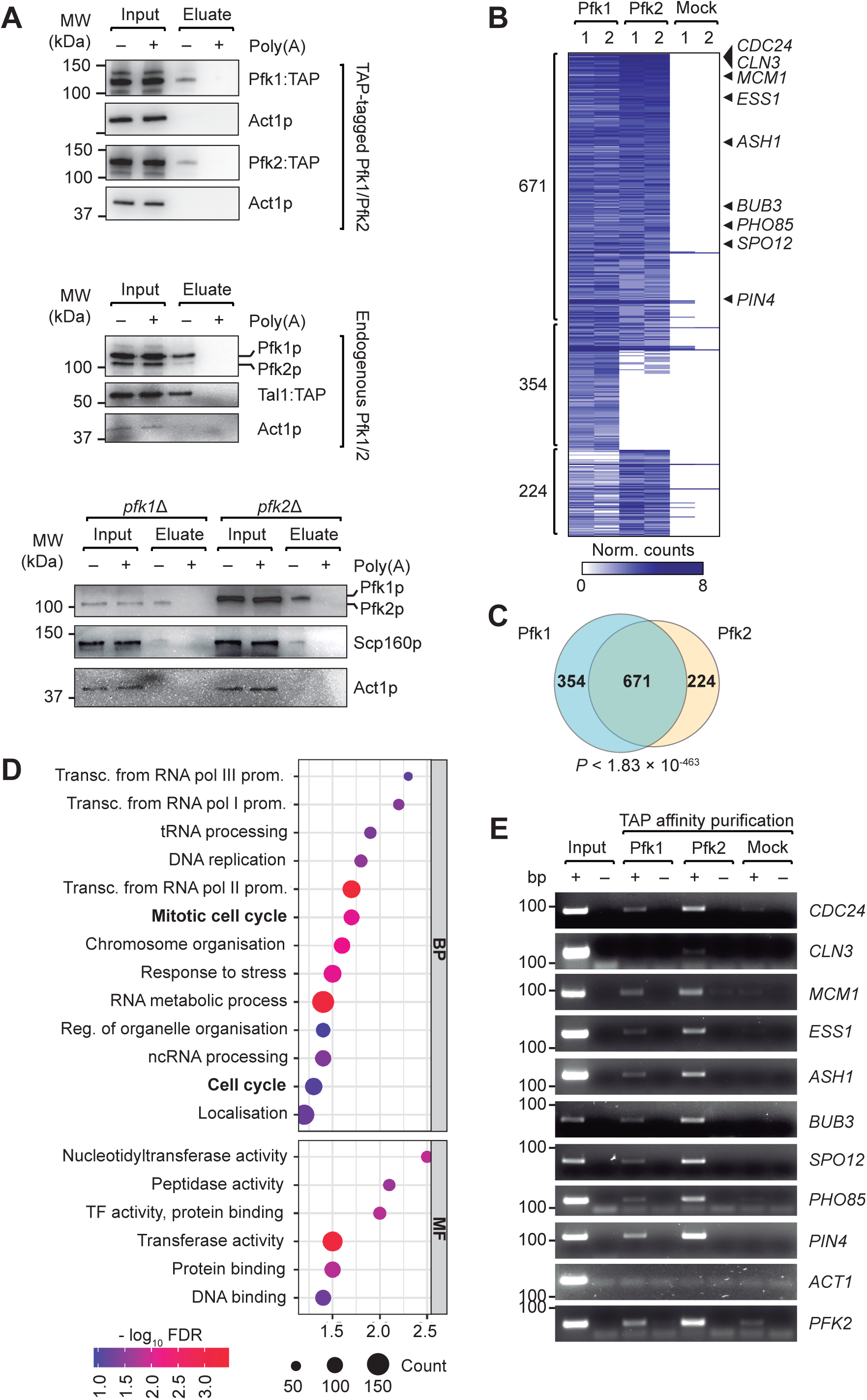
Pfk1p and Pfk2p bind to functionally related RNAs *in vivo*. (A) Immunoblot analysis of RIC eluates. Top: Pfk1:TAP and Pfk2:TAP detected with Peroxidase-Anti-Peroxidase Soluble Complex (PAP) reagent. Middle: endogenous Pfk1p and Pfk2 proteins detected with Pfk antibodies; Tal1:TAP is as non-conventional RBP ^5^. Bottom: Endogenous Pfk1p and Pfk2p detected in *pfk1Δ* and *pfk2Δ* cells. Scp160p is a RBP control; Act1p is non-RBP negative control. Input refers to cell extracts; poly(A) designates the addition of excess competitor polyadenylic acids. (B) Heatmap representation of the abundance of 1,249 fRIP selected Pfk1p and Pfk2 RNA targets. Columns refer to independent Pfk1:TAP, Pfk2:TAP, and untagged mock control fRIPs; rows denote individual transcripts. The white-blue colour bar represents normalised read counts for respective transcripts. (C) Venn diagram showing overlap of Pfk1p and Pfk2 mRNA targets. The *p*-value (hypergeometric test) relates to the significance of overlap. (D) Selection of GO terms overrepresented among 671 common Pfk1p and Pfk2 RNA targets. The diameter of the circle is proportional to the number of RNA targets, the colormap refers to the *-*log_10_ FDR. *X*-axis specifies the enrichment score. GO categories: BP, biological process; MF, molecular function. (E) Agarose gel showing products from RT-PCR reactions for detection of cell cycle related target mRNAs in fRIP eluates as marked in (B). Actin (*ACT1*) is a negative control. *PFK2* was previously shown in association with Pfk2p ^5^.

### Pfk1p and Pfk2p bind hundreds of functionally related mRNAs in cells

Next, we profiled the RNA targets for Pfk1p and Pfk2p by carrying out formaldehyde-assisted crosslinking and RNA-binding protein immunoprecipitation followed by high throughput sequencing of bound RNAs (fRIP-seq; see Methods). fRIP-seq was performed on yeast heterozygous diploid cells expressing *PFK1:TAP/PFK1*, *PFK2:TAP/PFK2* and an untagged parental wild-type strain that served as an IP background control to identify unspecific binding to the beads. To enable the estimation of relative enrichment scores, we also performed RNA-seq on ribosomal RNA depleted total RNA isolated from extracts. The IP efficiency was tested and confirmed that our chemical crosslinking does not interfere with the capture of the tagged proteins (Figure S1A).

Overall, 1,025 and 895 transcripts were significantly associated with Pfk1 and Pfk2, respectively (FDR ≤ 5%, ≥ 4-fold enriched over mock-IP control; Table S1A). Among those, 671 transcripts were common targets for both Pfk proteins (*p* = 1.83 × 10^-463^) (Figure 1B,C). Interestingly, RNA biotype analysis showed that most of the associated transcripts were mRNAs (1,017 and 867 of the Pfk1p and Pfk2p RNA targets, respectively; Figure S1B), which is different to other yeast glycolytic enzymes that preferentially interacted with short ncRNAs such as tRNAs ^21^. Nevertheless, 13 tRNAs were associated with Pfk2p including the glutamine-tRNA (tRNA-gln) *CDC65*, which has a role in the regulation of entry of cells into the cell cycle, while Pfk1p was only associated with one tRNA (tRNA-Ile) also bound by Pfk2p.

We next searched for significantly enriched Gene Ontology (GO) terms among Pfk1p and Pfk2p RNA targets. While no particular GO terms were enriched for RNA targets that were exclusive to either Pfk subunit, the 671 common RNA targets were significantly enriched (FDR < 0.05) for mRNAs coding for proteins acting in transcription (*e.g.,* ‘transcription from RNA polymerase II promoter’, *p* = 1.7 × 10^-5^), DNA replication (*p* = 0.007), chromatin organisation (*p* = 0.0022), and associated with the mitotic cell cycle (*p* = 7.4 × 10^-4^) (Figure 1D; Table S1B). Intrigued by this link to the cell cycle, we further validated the interaction of Pfk1 and Pfk2 with cell cycle factor mRNAs with independent fRIP followed by RT-PCR (Figure 1E). Notably, we observed stronger PCR signals with Pfk2 than with Pfk1 fRIPs, which could indicate preferential binding to Pfk2p. Overall, these findings indicate that Pfk1 and Pfk2 proteins interact with functionally related mRNAs which could facilitate the coordination of post-transcriptional gene regulation.

### Pkf2p binds short RNA sequence repeats *in vitro*

To identify common sequence/structural elements among mRNA targets that could mediate selective interactions with Pfk proteins, we retrieved the coding sequences (CDSs) as well as the 3’- and 5’-UTR sequences for Pfk1 and Pfk2 mRNA targets from the *Saccharomyces* Genome Database (SGD) and searched for the presence of common motifs using Multiple Expectation Maximization for Motif Elicitation (MEME) as an unbiased motif discovery tool (see Materials and Methods). UC-rich (UCRE) and GA-rich sequence elements (GARE) were overrepresented within the CDSs of RNA targets of Pfk1p and Pfk2p (Figure 2A, Table S2). Both motifs are arranged as trinucleotide repeats, spanning at least 4 repeats covering 11-12 nucleotides (nts). Interestingly, these triplet repeats are preferentially positioned in frame, (*i.e.,* 96% of the GARE and 87% of the UCRE repeats in Pfk2p mRNA targets), with GA(C/U) coding for the acidic amino acid aspartate (D), GA(A/G) coding for glutamate (E); and UCN coding for the polar uncharged amino acid serine (S). Hence, proteins encoded by those mRNAs bearing stretches of D/E amino acids are also overrepresented among Pfk’s mRNA targets (*e.g.,* MEME identified a stretch of eight D/E amino acids among Pfk2p mRNA target encoded proteins; 156 sites in 101 proteins, *p* = 3 × 10^-18^) (Table S2). Searching for the occurrence of UCRE and GARE motifs in all CDS of the yeast genome further confirmed the significant enrichment of these motifs among experimentally determined Pfk targets, reaffirming that these motifs could be selective recognition sites for Pfk proteins. However, the recorded in frame periodicity of the motif triplets is apparently reconciled across all CDSs of the genome (*i.e*., 7,973 (94%) of the 8,400 GARE motifs, and 3,504 (81.7%) of the 4,317 UCRE’s encoded all genomic CDS are in frame). Thus, the seen periodicity seems not selective for Pfk RNA associations (Table S2).

**Figure 2.**
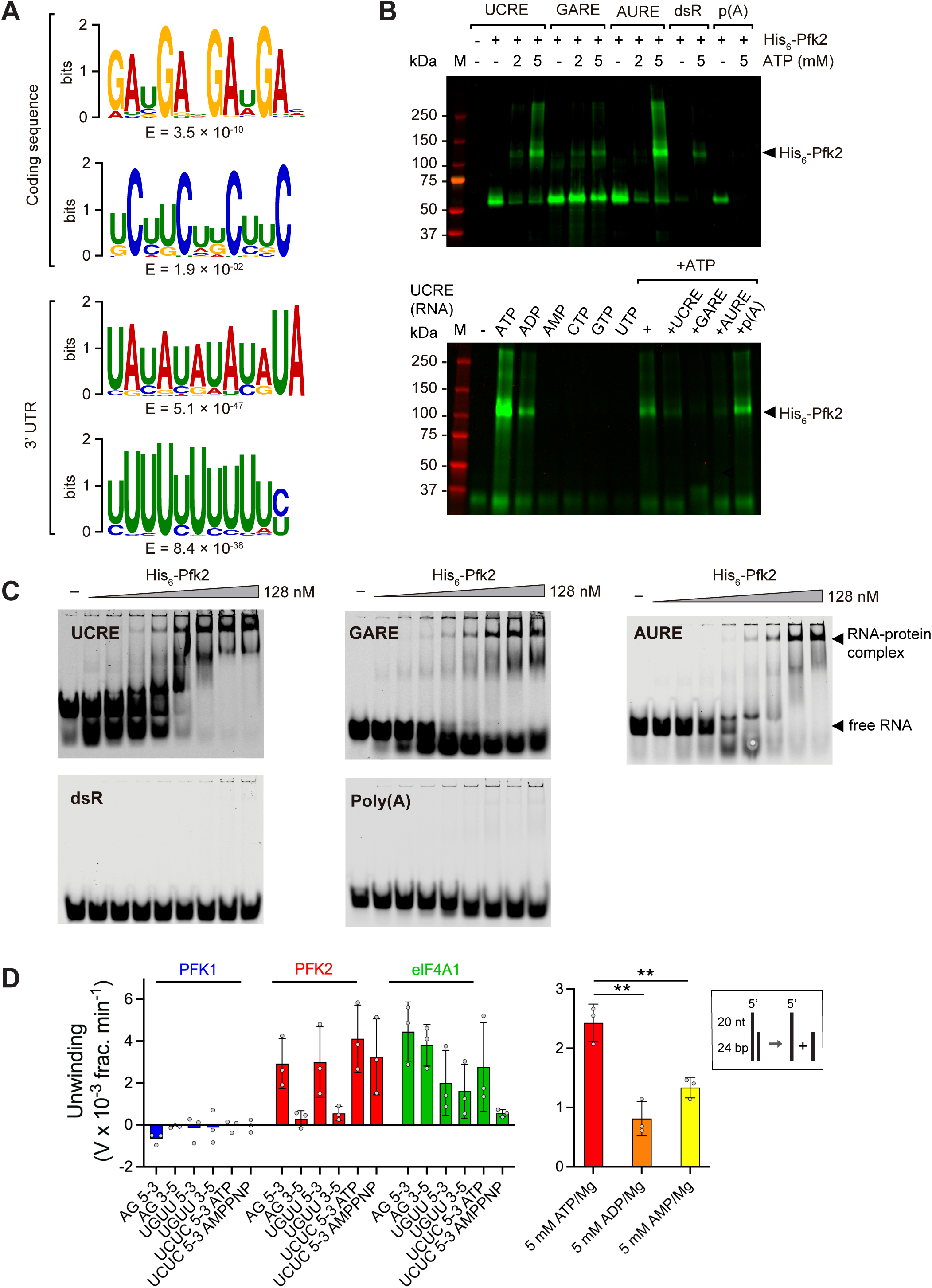
Pfk2p interacts with short RNA motifs and has directional RNA-unwinding activity *in vitro*. (A) Overrepresented sequence motifs in the ORF and 3’UTRs of Pfk2p RNA targets identified with MEME. E-values are indicated below each motif. (B) UV-crosslinking experiments with recombinant His_6_-Pfk2 and fluorescently labelled RNA oligos, including UCRE, GARE, and AURE RNA oligos matching the consensus sequences as well as double-stranded RNA motif (dsR) and poly(A) RNA (pA). The top gel illustrates dependence of interactions upon addition of Mg-ATP at indicated concentrations. The gel at the bottom shows interaction with UCRE RNA in the presence of 5 mM of indicated Mg-complexed nucleoside phosphates. A marker (M) with molecular weights in kDa is indicated to the left. (C) REMSAs with increasing concentration of His_6_-Pfk2p and the indicated RNA substrates in the presence of 5 mM Mg-ATP. Free RNA and the largest RNA-protein complex are marked to the far right. (D) RNA unwinding by Pfk1p, Pfk2p and eIF4A1 using 24 bp substrates with the indicated 20 nt repeat overhang sequences at the 5’ or 3’ ends. Data are mean values with standard deviations (± stdev) from repeat experiments (n=3). The UCUC 5-3 RNA tested with indicated adenosine phosphates is shown to the right (student’s *t*-test, \*\**p* < 0.01). A reaction scheme for 5’ overhang RNA substrates is boxed.

Likewise, motif searches with UTR sequences revealed enrichment of an AU-rich element (AURE) in 3’UTRs, as well as less-well defined A-rich and U-rich sequences in 5’- and 3’UTRs. The latter is reminiscent to a U-rich element previously identified in Pfk2p PAR-CLIP studies ^7^ (Figure 2A, Table S2). Importantly, the ‘U-rich’ elements contained interspersed ‘C’ nucleotides; and the ‘A-rich’ elements are intermingled with ‘G’ nucleotides which seem distantly related to the UCRE and GARE’s identified in CDS, respectively. A search with the individual consensus motifs across all genome-encoded UTR sequences revealed that the U-rich element located in 3’UTR, and the A-rich elements in the 5’UTR were both significantly enriched, while the AURE was not. However, this statistical analysis may be substantially obtruded by the limited information contained in A/U-rich sequences that are prevalent in the 3’UTR (*i.e*., 2,925 UAUAUAUAUAUA sites in 1,144 different 3’UTRs, as compared to 354 sites in 185 5’UTRs, and 319 sites in 137 different CDS (Table S2).

To test for direct interaction of Pfk proteins with those RNA elements, we then performed RNA-protein interaction assays with fluorescently labelled RNA oligos. Therefore, we designed IR800-labelled UCRE, GARE, AURE as well as poly(U) RNA oligos matching the consensus sequences, and we used labelled poly(A) RNA as a control.. Given that UCRE and GARE are complementary to each other and could form double-stranded RNA (dsRNA), we also designed an RNA oligo with complementary UCRE and GARE that could form a dsRNA structure (dsMotif/dsR). We then performed UV-crosslinking experiments and RNA electrophoretic mobility assays (REMSA) with recombinant His_6_-tagged Pfk1 and Pfk2 proteins expressed and purified from *E. coli* (Figure S2A). Since we got inconsistent data with Pfk1p regarding its RNA-binding specificity, we further focused our analysis on Pfk2p. UV-crosslinking experiments confirmed interactions of His_6_-Pfk2p to GARE, UCRE, AURE and poly(U) RNAs, which were competed by addition of excess unlabelled RNA (Figure 2B, Figure S2B). Surprisingly though, we found that interactions between RNA and full-length His6-Pfk2p were dependent on the presence of Mg-ATP. The addition of Mg-ADP slightly diminished RNA-protein interactions, while Mg-AMP and other nucleoside triphosphates (CTP, GTP, UTP) significantly reduced RNA binding of full-length His6-Pfk2p upon crosslinking. Noteworthy, in the absence of ATP, smaller-sized crosslinked products became apparent in the gel that varied in size and intensities between batches of purified recombinant proteins, possibly representing degradation products that were not further investigated. Finally, we performed REMSA in the presence of Mg-ATP to confirm interactions and to estimate binding affinities. In agreement with UV-crosslinking experiments, strong interaction of Pfk2p with UCRE, GARE and AURE in the nanomolar range was observed. Thereby, it appeared that Pfk2p could form different high-molecular complexes on the RNA not seen with dsMotif and poly(A) control RNA oligos under our chosen experimental conditions. Considering the largest protein-RNA complex, strongest interactions were estimated with UCRE (K_d_ ∼ 13 nM) and some weaker association with AURE (Kd ∼ 29 nM) and the GARE (Kd ∼ 40 nM) (Figure 2C). In conclusion, our data show that His_6_-Pfk2p can selectively interact with short GA-, UC- and AU/U-rich sequences in an Mg-ATP-dependent manner with high affinity.

### Pfk2p has RNA unwinding activity

Curious about the appearance of various intermediate products in REMSA experiments (Figure 2C), which might suggest conformational changes in RNA or RNA-protein complexes, we considered Pfk2p could drive structural modifications in the RNA. We therefore performed real-time fluorescence-based RNA unwinding assays using RNA substrates previously tested with eIF1A1, a human DEAD-box family RNA helicase involved in translation initiation ^22^. The RNA substrates contained an identical 24 bp duplex with 20 nt overhang sequences related to our identified GA and UC-rich sequence elements placed either at the 5’ or the 3’ end of the RNA ^22^. Surprisingly, we found that His_6_-Pfk2p displayed RNA unwinding activity with RNA substrates bearing the overhang sequence at the 5’ end, while no unwinding activity was detected with overhang sequences placed at the 3’ end (Figure 2D). As expected, eIF1A1 displayed unwinding activity irrespective of the position of the overhangs. No activity in either direction was detectable with likewise expressed and purified His_6_-Pfk1p. These results suggest that Pfk2p but not Pfk1p has 5’ to 3’ directional unwinding activity applied to short RNA duplexes. Furthermore, while eIF4A1 RNA unwinding activity was abrogated in the presence of 5 mM adenylyl-imidodiphosphate (AMPPNP, a non-hydrolysable ATP analog), His_6_-Pfk2p remained similarly active. Hence, ATP hydrolysis may not be strictly required for Pfk2p’s unwinding activity. Interestingly, the addition of respective diphosphates and monophosphates (ADP and AMP) reduced Pfk2p’s unwinding activity by 40-60% (Figure 2D). While this inhibition is reminiscent to the previously observed reduced UV crosslinking efficiency of full-length Pfk2p to RNA in the presence of ADP and AMP, residual unwinding activity may be attributed to competition with ATP still bound to Pfk2p from the purification procedure.

### Pfk RNA and ribosome interactions are dependent on the energy-status of cells

Since Pfk2p-RNA interactions are ATP-dependent *in vitro*, we next wondered whether the association with RNA could depend on the energy status of cells *in vivo*. Hence, we performed a RIC experiment with cells subjected to glucose depletion and after re-addition of glucose (Figure 3A). The removal of glucose from the media immediately affects the levels of various metabolites and drastically depletes cellular ATP levels by ∼70% within minutes while increasing the cellular AMP/ATP ratio in budding yeast ^23–25^. As expected, we found that glucose starvation abolished Pfk1:TAP and Pfk2:TAP binding to poly(A) RNA and re-addition of glucose to media largely restored their interactions with RNA (Figure 3B). This change in RNA association was not observed with Pgk1:TAP, which is another RNA-binding glycolytic enzyme that remained bound to poly(A) RNA under all conditions (Figure 3B). Essentially, in agreement with *in vitro* RNA binding assays, these results suggest that interactions of Pfk subunits with RNA in cells are ‘energy dependent’, consistent with the lower availability of cellular ATP in glucose starved cells.

**Figure 3.**
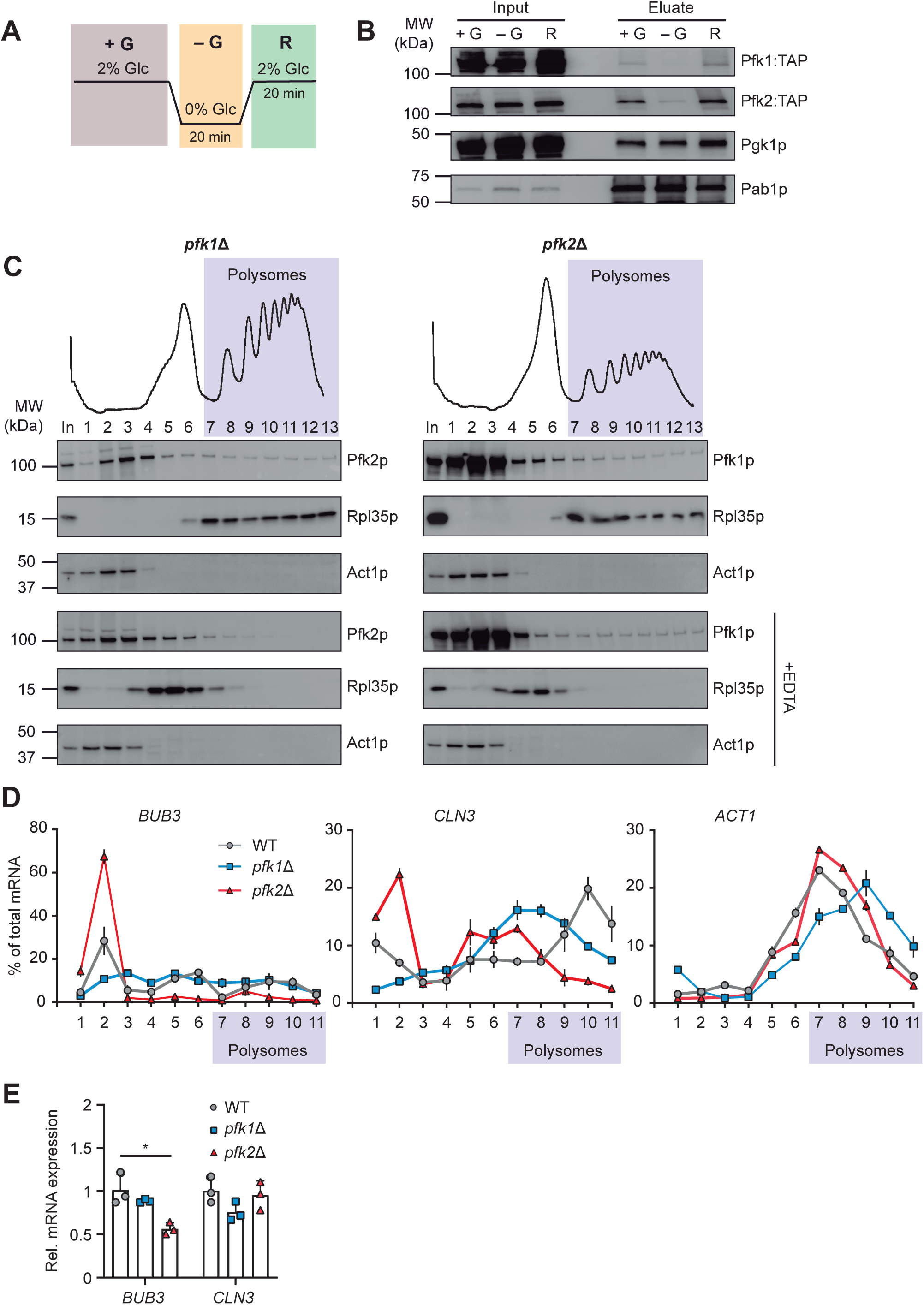
Pfk2p dynamically associates with mRNAs and polysomes and impacts translation. (A) Experimental design for glucose (glc) recovery experiments. Cells were grown in YPD (+G), then starved in media lacking glc for 20 min (-G) and recovered upon re-addition of glc for 20 min (R). (B) Immunoblot of input extracts (left) and RIC eluates (right) from glc recovery experiments at indicated stages (+G, -G, R). Monitored proteins are labelled to the right; a molecular weight marker is indicated to the left. (C) Polysomal absorbance profiles of *pfk1Δ* and *pfk2Δ* cells grown in YPD medium are shown at the top. Fractions numbers are indicated and those containing polysomes are highlighted in purple. Immunoblot analysis of fractions monitoring the distribution of Pfk1p and Pfk2p; and upon treatment of extracts with 30 mM EDTA to dissociate ribosomes (see Methods; Figure S3) are given below. Rpl35p is a ribosomal protein of the large subunit, Act1p is a non-ribosomal associated control protein. (D) Distribution of *CLN3*, *BUB3* and *ACT1* mRNA levels across sub-polysomal and polysomal fractions (purple) obtained from wild-type (wt; grey), *pfk1Δ* (blue) and *pfk2Δ* (red) cells. RNA was isolated from each fraction and quantified by RT-qPCR. The *y*-axis denotes mRNA levels in each fraction calculated as the percentage of the total (mean values ± stdev, n=3). (E) *BUB3*, and *CLN3* mRNA levels relative to *ACT1* levels determined by RT-qPCR in total RNA isolated from wt, *pfk1Δ* and *pfk2Δ* cells (mean values ± stdev, n=3). \**p* < 0.05 (student’s *t*-test).

Since yeast Pfk2p and human orthologs such as the liver and platelet isoforms PFKL and PFKP have previously been detected in polysomes and in association with cytoplasmic ribosomes, respectively ^26,27^, we wondered whether the yeast Pfk paralogous could assist in translation. Hence, we performed sucrose density fractionations with wild type, *pfk1*Δ and *pfk2*Δ strains to test if Pfk subunits are associated with polysomes. While *pfk1Δ* cells showed similar polysomal profiles like wild-type cells (ratio of polysomes to subpolysomes (P/S) = 1.85 – 1.95), *pfk2*Δ cells showed a ∼50% decrease in polysomes (P/S = 0.85) indicating diminished global translation (Figure 3C, Figure S3). This is in line with the reported slow-growth of *pfk2*Δ cells, which manifests in reduced protein synthesis. We further found that ∼20% of Pfk1p and ∼11% Pfk2p co-sedimented with polysomes in heavy sucrose fractions. However, only Pfk2p dissociated from these polysomal fractions upon treatment of extracts with 30 mM EDTA, a condition that disassembles polysomes and ribosomes.

In contrast Pfk1p did not disassemble, suggesting that Pfk1p could be part of another high order complex the co-sediments with polysomes, such as previously reported highly filamentous structures that can form in the absence of *PFK2* ^28^. We next tested whether the association of Pfk2p with ribosomes is affected by glucose levels in the growth media as previously monitored in RIC experiments (Figure 3A). We found that Pfk2p dissociated from polysomes in *pfk1*Δ cells upon glucose removal and fully reassociated with polysomes after re-addition of glucose to media (Figure S3). Conversely, Pfk1p did not dissociate from high-density gradient fractions upon glucose removal and even after disassembly of polysomes with EDTA, corroborating the notion that Pfk1p is not directly associated with translating ribosomes in the absence of Pfk2p (Figure S3). Taken together, these results suggest dynamic and reversible association of Pfk2p - but not of Pfk1p - with translating ribosomes.

As many ribosome-associated RBPs act as translational regulators on selected mRNAs, we further monitored the distribution of Pfk mRNA targets, namely *cyclin 3* (*CLN3*) and budding uninhibited benzimidazole 3 (*BUB3*) coding for proteins involved in cell cycle regulation and DNA replication, across the sucrose density fractions isolated from wild-type, *pfk1*Δ and *pfk2*Δ cells with RT-qPCR. Indeed, we found a dramatic shift of both *BUB3 and CLN3* mRNAs from heavy polysome fractions to ‘subpolysomal’ fractions (comprising RNPs, free ribosomal subunits and monosomes) in *pfk2*Δ but not in *pfk1*Δ cells and as compared to wild-type cells (*BUB3*; ∼10% in polysomal fractions of *pfk2Δ* cells compared to 40% and 32% in *pfk1Δ* and wt cells, respectively; *CLN3*, 31% in polysomal fraction of *pfk2Δ,* 63% and 60% in *pfk1Δ* and wt cells, respectively). No change was seen in the distribution of *ACT1* mRNA as a non-target control (Figure 3D). The corresponding total RNA levels isolated from extracts were not altered for *CLN3* in *pfk1*Δ and *pfk2*Δ compared to wild-type cells; however, *BUB3* mRNA levels were reduced in *pfk2Δ* cells, which could indicate reduced mRNA stability associated with the observed drastic shift to ribosome devoid fractions (Figure 3E). Hence, the dynamic association of Pfk2p with polysomes along the severely impacted co-sedimentation of selected Pfk target mRNAs with polysomes in *pfk2Δ* cells, suggests that Pfk2p could act as a translational activator for selected mRNAs.

### Deletion of *PFK2* affects abundance of a large fraction of the proteome

To assess the proteome-wide impact of PFK expression and the possible implications for cell-physiology, we next profiled changes of the of the proteome of *pfk1*Δ and *pfk2*Δ mutants compared to wild-type cells using isobaric tandem mass tag (TMT) based quantitative proteomics (see Methods). We also profiled slow-growing *map1*Δ cells ^29,30^ to evaluate whether altered protein levels could simply be associated with slow growth as reported for *pfk2*Δ strains. Data were obtained for 3,856 proteins (n=3; 1% FDR). Irrespective of the chosen cut-off, the largest fraction of protein level changes was seen in *pfk2*Δ cells (*e.g*., 1,740 proteins at FDR ≤ 10%). Substantial fractions also changed in *map1Δ* cells (1,100 proteins, FDR ≤ 10%), whereas considerably less changes were observed in *pfk1*Δ cells (672 proteins, FDR ≤ 10%)(Figure 4A; MS data given in Table S3). As may be expected, proteins changing in *pfk1Δ* and *pfk2Δ* cells greatly overlapped, most of them showing reduced relative abundances (Figure 4A). GO analysis revealed that the commonly ‘down-regulated’ proteins, *i.e.,* less abundant proteins in *pfk1Δ, pfk2Δ* and *map1Δ* compared to wild-type cells, refer to ‘cytoplasmic translation’, ‘ribosome biogenesis’ and/or are part of ‘ribonucleoprotein complexes’ (Figure 4B, complete GO analysis given in Table S4). Along the commonly observed slight increased levels of proteins associated with the oxidative stress response, these changes are obviously not selective for the lack of *PFK* genes as also seen in slow-growing *map1Δ* cells. A second subgroup of proteins showing increased abundances in *pfk1*Δ and *pfk2*Δ but not in *map1Δ* cells comprised components of the ‘carbon metabolic process’ and the ‘generation of precursor metabolites and energy’ including glycolytic enzymes. The increased levels of carbon metabolism components in *pfk* mutants could relate to a compensation mechanism to cope with the reduced glycolytic activity in *pfk* mutants. Finally, a third group of proteins acting in DNA replication, tRNA processing and transcription were particularly lower abundant in *pfk2*Δ but not in *pfk1*Δ cells, i.e. representing *pfk2Δ* specific implications in the proteome (Figure 4B, Table S4). These functional themes were reminiscent to the relationships of experimentally determined Pfk mRNA targets and hence, indicated functional links between mRNA targets and aberrant expression in *pfk2Δ* cells.

**Figure 4.**
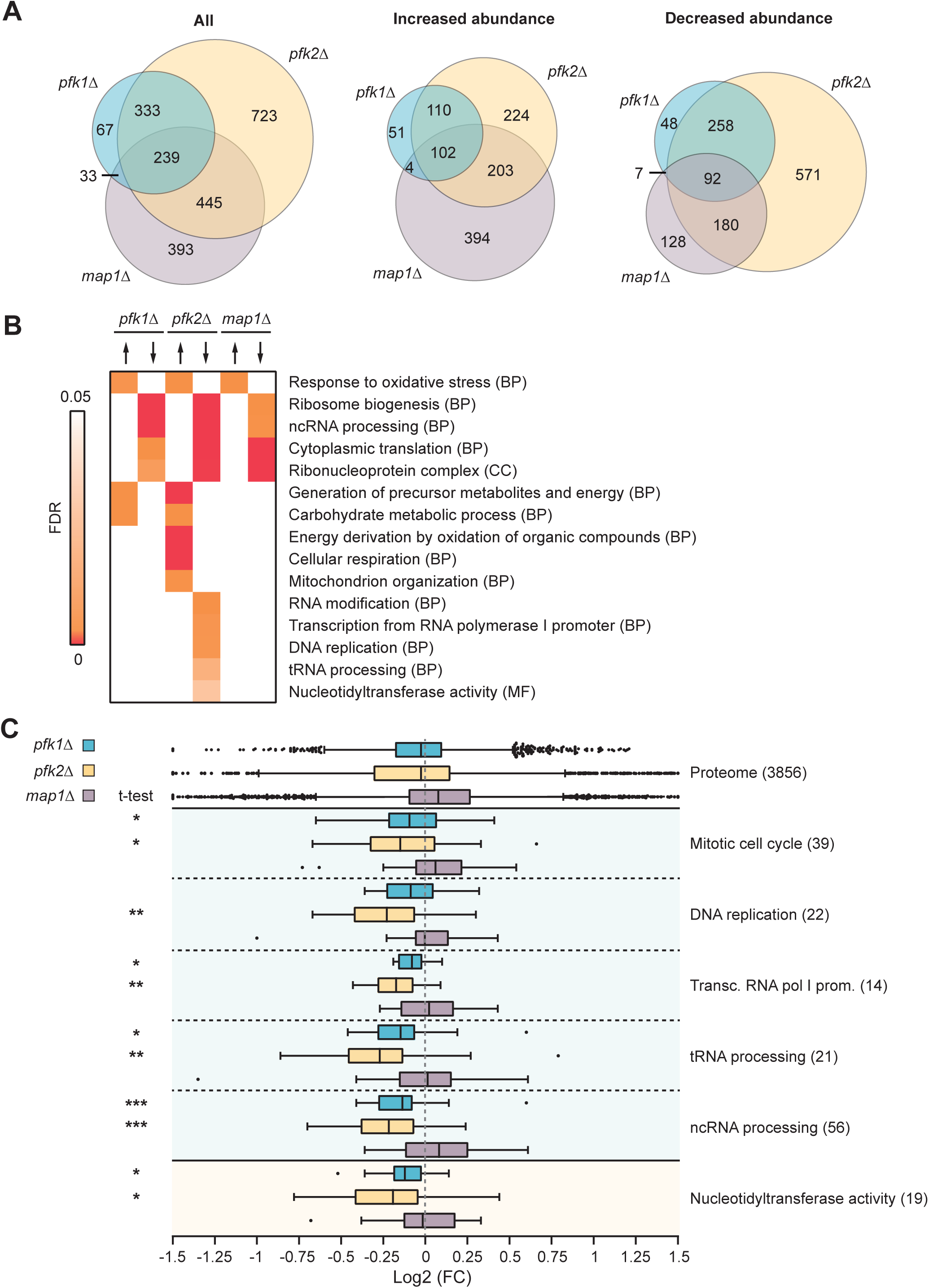
Proteome changes in *pfk1*Δ, *pfk2*Δ and *map1*Δ cells. (A) Venn Diagram displaying the overlap of numbers of proteins with altered levels (FDR ≤ 10%) in the mutant compared to wild-type (wt) cells. (B) Heatmap displaying a subset of GO terms (rows) enriched among the proteins with increased (upright arrow) or reduced (downward arrow) levels in indicated mutants (columns). The colour intensity corresponds to Bonferroni corrected FDRs. (C) Boxplots depicting relative fold-changes of protein levels in *pfk1Δ* (blue), *pfk2Δ* (yellow) and *map1Δ* (grey) cells compared to wt cells (log_2_ scale). Whiskers extend from the 10^th^ to the 90^th^ percentile. The distribution of all proteins is shown at the top. GO terms are indicated to the left and the number of plotted Pfk2p mRNA target encoded proteins within the respective GO group is indicated in brackets. Asterisks refer to *p*-values determined in a student’s *t*-test with Welch’s correction comparing the distribution of Pfk2p mRNA target encoded proteins assigned to specified GO term with the distribution of all measured features: \*\*\**p* < 0.001; \*\**p* < 0.01; \**p* < 0.05.

To further establish proteome-wide connections to mRNA targets, we mapped experimentally determined Pfk1p and Pfk2p targets to changes in proteomes (MS data was available for 730 and 620 of Pfk1p and Pfk2p mRNA targets, respectively). We found only a slight overrepresentation of proteins encoded by Pfk1p and Pfk2p mRNA targets among the proteins selected with 10% FDR in *pfk1*Δ and *pfk2*Δ cells, respectively (Pfk1p: 111 proteins, *p* = 0.043; Pfk2p: 274 proteins, *p* = 0.3). Of note, choosing a more stringent FDRs, significant associations were also revealed for Pfk2p mRNA target encoded proteins (FDR < 1%, 88 proteins, *p* = 0.051; among those 41 less abundant in mutants, *p* = 0.002). However, combinatorial control through other RBPs and secondary effects induced through metabolic changes in *pfk* mutants may confuse ‘simple’ associations between changes in protein levels and the encoded mRNA targets. Hence, we searched for specific instances on subsets of mRNAs attributed to particular GO terms. Indeed, we found significant directional responses at the protein levels for functionally related mRNA subsets (Figure 4C). For instance, reduced protein levels for Pfk2p mRNA targets coding for the ‘mitotic cell cycle’, ‘DNA replication’ and Pol I related subjects like ‘tRNA processing’ were found. Importantly, the generally decreased abundance of those functionally-related target groups, which was slightly more prevalent in *pfk2Δ* than in *pfk1Δ* cells, agrees with Pfk2’s proposed role as a translational activator.

### *PFK2* modulates cell cycle progression independent of its catalytic activity

We wondered whether the observed reduced expression of cell-cycle related mRNAs is also mirrored by the cell’s phenotype. *Pkf2Δ* cells elicit a slow-growth phenotype not seen with *pkf1Δ cells* but the source of these phenotypic differences remained unclear ^20^. Furthermore, it was noted that *in vitro* measured catalytic activity is depleted in cell extracts derived from either *pfk* gene knock-out strains, indicating that catalytic activity is unlikely a major source driving the *pfk2Δ* phenotype ^20^. To confirm these findings in our yeast strains, we first compared the growth of wild-type, *pfk1*Δ and *pfk2*Δ cells, and reintroduced wild-type genes as well as mutants with compromised catalytic activity for functional complementation (Figure 5A). As expected, *pfk2Δ* cells showed a slow-growth phenotype while *pfk1*Δ cells grew like wild-type cells, confirming that the presence of heteroctameric Pfk1p-Pfk2p complexes is not required for cell fitness. Furthermore, the slow-growth of *pfk2Δ* cells was rescued by reintroduction of the wild-type *PFK2* gene as well as catalytic mutants expressed form a single-copy plasmid. Specifically, an exchange of aspartate 348 with serine (D348S) that eliminates the proton acceptor in the substrate binding site reduced activity up to 50%. Likewise, mutation of aspartate 301 to threonine (D301T), which involves the catalytic Mg^2+-^ATP binding site reduced catalytic activity by almost 40% compared to *PFK2* rescued cells, which agrees with previous observations ^19^ (Figure 5B). To see whether both mutations could act synergistically, we further created a double mutant (DM) at both sites (D301T, D348S). The DM showed reduced enzymatic activity and rescued the growth defect of *pfk2Δ* cells like single catalytic mutants and the wild-type *PFK2* gene. This suggests that remaining catalytic activity is contributed by the residual Pfk1p subunits in complex with the catalytically dead Pfk2p (since neither extract from *pfk1Δ* nor pfk2Δ cells have measurable enzymatic activity *in vitro*). Conversely, the slow-growth phenotype of *pfk2Δ* cells can obviously not be explained by the reduced catalytic activity of Pfk2p.

**Figure 5.**
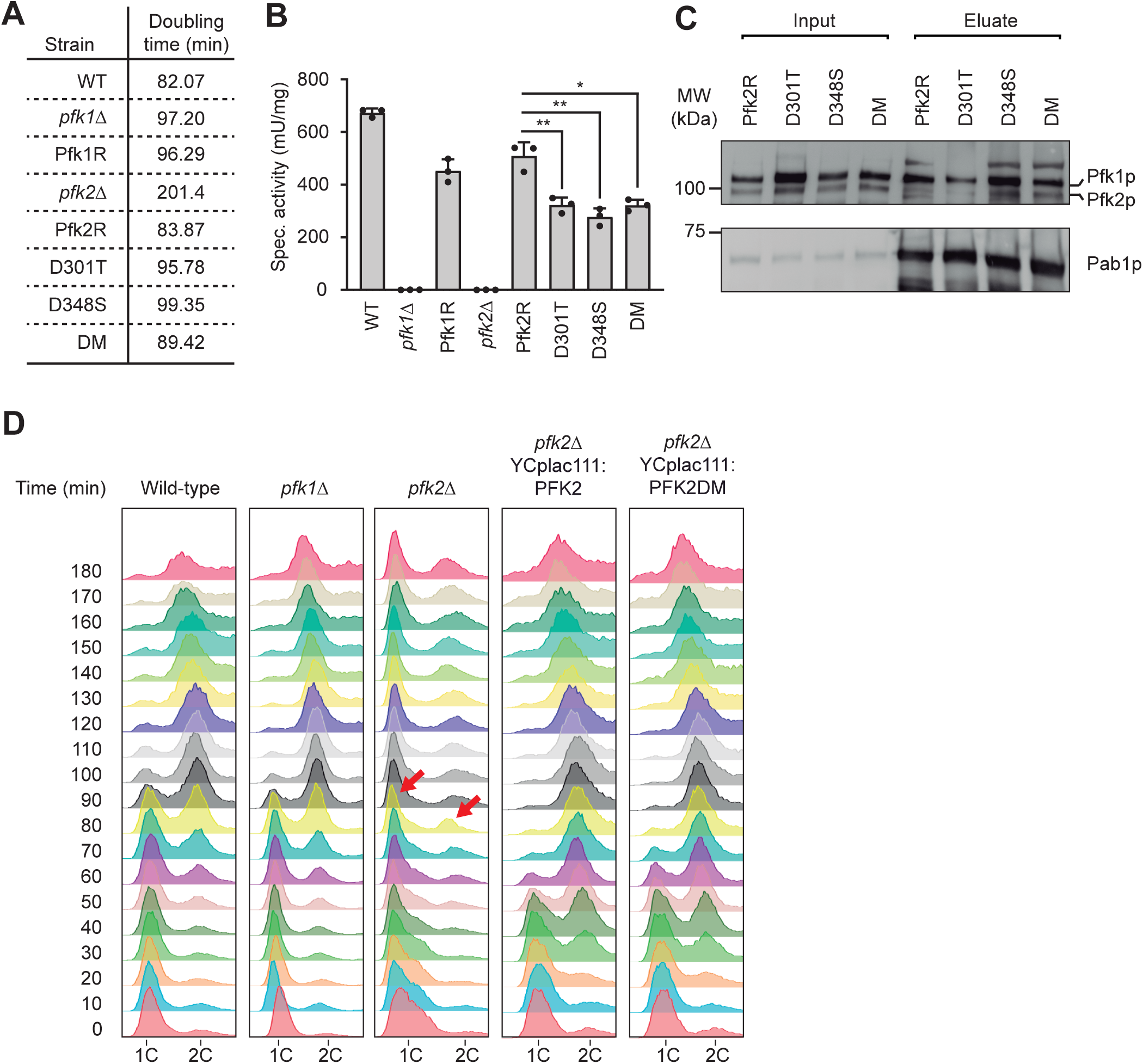
Slow growth and cell cycle progression defects of *pfk2Δ* cells can be rescued with catalytic mutants. (A) Doubling times of indicated yeast strains grown in YPD media at 30 °C. (B) PFK activity measurements. Bar depicts PFK specific activity (mU/mg) of 5-6 paired replicate measurements in extracts derived from indicated cells. Standard deviations are indicated in brackets; student’s *t*-test, \*\**p* < 0.01; \**p* < 0.05. (C) Immunoblot of RIC eluates for indicated strains. The input extract is shown to the left. (D) Analysis of cell cycle progression. Yeast strains indicated at the top were synchronized in G1 phase using α-factor. After release from α-factor, samples were collected every 10 min (*y*-axis), fixed, and analysed for DNA content (1C/ 2C; *x*-axis) at specified time points with flow cytometry. Red arrows depict aberrant ‘1C’ and ‘2C’ peaks in *pfk2Δ* cells.

To evaluate potential impact on the association of the Pfk complex with RNA, we performed RIC, revealing that complementation with single and double catalytically dead *pfk2* mutants did not greatly affect poly(A) RNA association *in vivo* (Figure 5C). Thus, unlike enzymatic activity, the formation of an RNA-binding complex is not significantly compromised in those mutants, which can rescue the slow-growth phenotype (Figure 5A).

Finally, we assessed potential cell-cycle defects within the single deletion mutants. Whilst *pfk1*Δ mutants exhibited cell cycle progression like wild-type cells, we observed that a significant portion of *pfk2*Δ cells showed delayed progression from G1 into S phase and further propagation into G2/M phase (Figure 5D). Importantly, the cell cycle defect was fully complemented by reintroducing either exogenously expressed *PFK2* or the catalytic double mutant (DM). Hence, as described above, the cell cycle defect in *pfk2*Δ cells can neither be explained by the absence of heteromeric complexes nor the compromised enzymatic activity of Pfk2. Accordingly, we postulate that the absence of Pfk2p associated RNA binding functions along with reduced translation of cell cycle regulators contributes to the observed phenotype.

## Discussion

Despite being studied for over 70 years for its role in glycolysis, only recent evidence suggested that phosphofructokinase (PFK) also binds RNA, although the exact role of this function has remained unclear. In this study, we gained insights into yeast PFK RNA binding and discovered a first functional distinction between the two paralogous subunits, Pfk1p and Pfk2p. Specifically, we demonstrate that Pfk2p acts as an ATP-dependent mRNA-binding protein that bears directional 5’-3’ dsRNA unwinding activity. Pfk2p enhances the expression of cell-cycle-related mRNAs at the post-transcriptional level, and it promotes cell cycle progression from G1 to S phase independent of the enzyme’s catalytic activity. Hence, we suggest that Pfk2p acts as a cellular control point or ‘molecular switch’. By doing so, it facilitates direct communication between the metabolic state and translation of cell cycle components to coordinate cell proliferation, enabling rapid and sensible responses to frequent nutritional changes experienced by unicellular organisms.

### Pfk2 acts almost like a canonical mRNA binding protein

We used a chemical crosslinking approach to identify cellular RNA targets for both Pfk1p and Pfk2p. This analysis revealed that both Pfk proteins bind to a substantially overlapping set of hundreds of different mRNAs and a few non-coding RNAs. Given that Pfk1 and Pfk2 form a stable hetero-octameric protein complex, this significant overlap in RNA targets may be expected; however, alternate multi- and monomeric complexes may also exist, as Pfk proteins can associate with cellular RNA independently of their paralog (Figure 1A). The commonly bound mRNAs encoded functionally related protein groups, a characteristic of canonical RBPs that form post-transcriptional operons or RNA regulons for coordination of post-transcriptional events ^31,32^. These functional associations - particularly those linked to the mitotic cell cycle - suggested roles for Pfk proteins in cell cycle control reminiscent to previous observations ^33^, which we finally confirmed (Figure 5D). Furthermore, consistent with canonical RBPs that recognize RNA via selective sequences or structural motifs, bioinformatic analysis identified short and unstructured single-stranded RNA sequences (UC-, GA-, AU-, or U-rich) that interact with Pfk2p with high affinity in the presence of Mg-ATP (Figure 2B,C). This finding is remarkable insofar as RBPs without canonical RBDs are rarely sequence-specific ^34^. Moreover, to our knowledge, Pfk2p is the first example of an unconventional RBP whose RNA interaction is ATP-dependent, as shown by *in vitro* binding assays and inferred *in vivo* through glucose removal, which rapidly depletes cellular ATP levels (Figure 3A,B). Therefore, the connection between bound mRNA identities and allocated phenotype along with distinct RNA sequences that can drive direct interactions suggests that Pfk2p acts similar to classical RBPs being an integral part of the yeast’s post-transcriptional regulatory system. As RNA-binding was reported for PFK orthologues across diverse organism and human cells ^1,8,35^, it will be interesting to explore insofar RNA-binding specificity and functions are evolutionary conserved.

### Pfk2p has non-conventional RNA unwinding activity

A prominent class of conserved enzymes that use ATP to bind or remodel RNA or ribonucleoprotein complexes (RNPs) are RNA helicases, which play diverse roles across RNA-related processes, from splicing to translation and degradation ^36,37^. Surprisingly, we discovered that recombinant Pfk2p, but not Pfk1p, demonstrated RNA unwinding activity with pronounced 5’ to 3’ directionality. This contrasts with the DEAD-box RNA helicase eIF4A, a member of the SF2 subfamily, which exhibits only limited strand displacement directionality (Figure 2D). In this regard, Pfk2p’s activity could resemble eukaryotic RNA helicases of the SF1b family, such as MOV10 with 5’ to 3’ directionality for strand displacement ^38^. However, the molecular mechanisms for strand-displacement by Pfk2p needs further investigation and remains speculative. For instance, analogous to other eukaryotic RNA helicases, which bear two RecA subdomains that close around the RNA upon NTP binding and promote strand-displacement, a conformational shift of Pfk2p induced by Mg-ATP could prompt the low-activity T-state which is known to come along with a re-orientation of the N-half (containing the active site) and C-terminal half (housing allosteric regulatory sites) in enzyme subunits ^9,39^. This could eventually facilitate RNA interaction and directional unwinding, which may involve Pfk2p oligomerization on the RNA. In this context, it would be interesting to see whether the N-terminally located glyoxylate I-like domain or intrinsically disordered regions (IDR) located nearby and thought to promote Pfk2p assembly into G-bodies are involved in guiding the observed directionality ^40^. Finally, ATP hydrolysis might be required for Pfk2p’s release from RNA substrates, stimulating a conformational shift to the high-activity R-state with bound ADP/AMP. Although speculative, this model could explain the retained RNA unwinding activity in the presence of a non-hydrolysable ATP analogue (AMPPNP) and the reduced activity observed upon competition with ADP or AMP, potentially shifting Pfk2p towards the high-activity R-state (Figure 2D).

### Pfk2p can promote the translation of cell cycle mRNAs

As observed for many RNA helicases, we found that translation is disrupted in *pfk2Δ* cells, further suggesting that Pfk2p specifically enhances the translation mRNAs coding for cell cycle regulators, such as *CLN3* and *BUB2* (Figure 3D). Notably, these translational defects occur exclusively in *pfk2Δ* cells. This correlates well with the observation that only Pfk2p, and not Pfk1p, associates with ribosomes in a manner dependent on the cell’s energy state, as found in the glucose-depletion assays (Figure S3). Since Pfk2p demonstrates RNA unwinding activity and associates with ribosomes, it seems likely that Pfk2p facilitates translation by resolving mRNA secondary structures or displacing mRNP complexes, thereby improving ribosome processivity and enhancing translation efficiency.

Proteomic analysis of *pfk1Δ*, *pfk2Δ* and *map1Δ* yeast mutants revealed that *pfk2Δ* exhibited the most significant changes on protein levels, with a substantial overlap of all three mutants among ‘downregulated’ proteins, particularly those involved in cytoplasmic translation and ribosome biogenesis (Figure 4A,B). Likewise, proteins related to oxidative stress showed increased levels in all mutants, indicating a generalized stress response. In contrast, proteins with roles in carbon metabolism were exclusively ‘upregulated’ in both *pfk1Δ* and *pfk2Δ*, likely compensating for reduced glycolytic activity. Moreover, proteins related to DNA replication and transcription were preferentially downregulated in *pfk2Δ*, suggesting a subunit-specific impact on the proteome (Figure 4B). Further analysis connected Pfk1p and Pfk2p mRNA targets to these proteomic changes, particularly in *pfk2Δ*. Specifically, the analysis of functional GO clusters revealed a significantly lower abundance of proteins encoded by Pfk2p mRNA targets associated with the mitotic cell cycle, DNA replication, and tRNA processing, supporting a role of Pfk2p in translational activation (Figure 4C). Overall, these results affirm Pfk2p’s specific impact on mRNA target expression and align with its role in cell cycle progression.

### Physiological relevance of Pfk2p mRNA binding

Phenotypically, *pfk2Δ* but not pfk1Δ cells, exhibit a pronounced slow-growth phenotype that can be effectively rescued by reintroducing exogenously expressed *PFK2* or its catalytically dead mutant alleles on a centromeric vector (Figure 5A). Consistent with earlier reports showing no detectable catalytic activity in extracts from either single mutant ^18,20,41^, our findings strongly suggest that the growth phenotype in *pfk2Δ* cells is not dependent on the enzyme’s catalytic functions (Figure 5B). Likewise, we observed that *pfk2Δ,* but not *pfk1Δ,* cells showed significantly delayed progression from the G1 to the S phase of the cell cycle independent of the enzyme’s catalytic activity (Figure 5D). Combined with evidence that Pfk2p binds to mRNAs coding for cell cycle-related proteins – and RNA binding remaining intact in catalytically inactive mutants (Figure 5B) - alongside Pfk2p’s role in translation and reduced levels of cell cycle proteins in *pfk2Δ* cells, we propose that the cell cycle defect arises from Pfk2p’s function as a post-transcriptional regulator of cell cycle genes.

### Is Pfk2p a molecular relay switch?

We further propose that Pfk2p acts as a molecular relay for transduction of metabolic signals to the post-transcriptional level to control cell cycle progression and possibly other processes (Figure 6). When cellular energy levels are low, Pfk proteins preferentially adopt the high activity state (R-state) through binding of AMP/ADP at allosteric activator sites and engage in glycolysis for ATP production but not binding to RNA. Conversely, when energy levels become sufficiently high, Pfk2p can adopt the low-activity conformation (T-state) with low glycolytic activity but binding to mRNAs. Through Pfk2p’s association with ribosomes, the translation of cell cycle and other transcripts is facilitated - possibly through Pfk2p’s RNA unwinding activity – which ultimately promotes cell cycle progression. In this way, Pfk2p could constitute a ‘molecular relay switch’ to balance cellular needs: cell proliferation is repressed in energy deprived cells where resources must be allocated to energy production to secure cell survival, while favourable conditions mounted by high energy levels reallocate resources towards production of proteins for cell proliferation and enabling colony expansion. Such a molecular switch at the post-transcriptional level can be highly efficient and rapid, allowing for immediate response to changes in available nutrients and circumvents the need for reshaping the transcriptome. As such, it creates an ‘economic’ solution for short-term adaptation to changing energy levels imposed by the availability of nutrients coordinating glycolytic flux and cell proliferation. Given that numerous metabolic enzymes have been shown to interact with RNA, potentially affecting both the bound RNAs and the enzymes themselves, it is plausible that many more relay switches between intermediary metabolites and RNA may exist through specialization of paralogous enzymes ^42^, forming a vast network that we are only beginning to explore.

**Figure 6.**
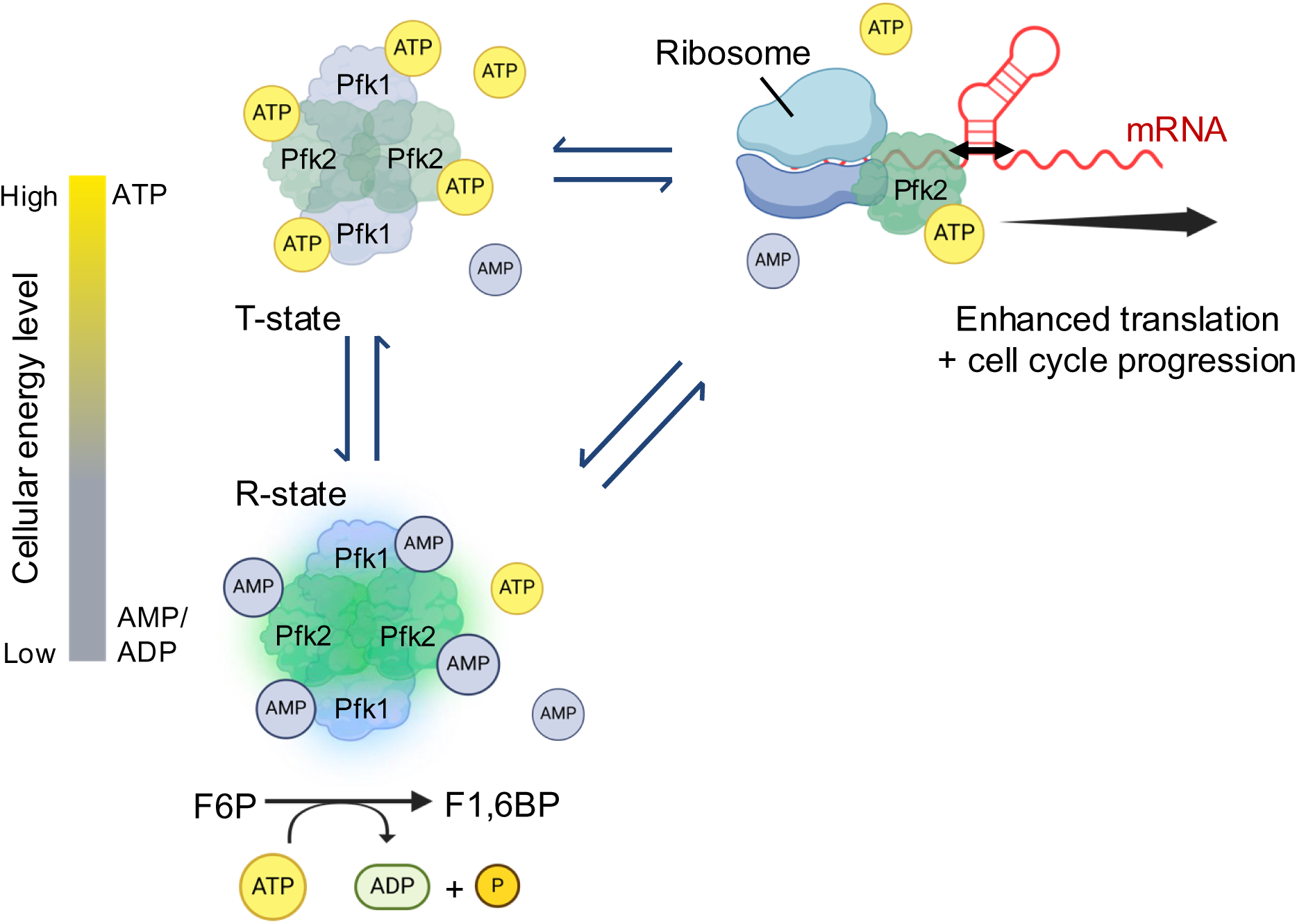
A relay switch model for yeast Pfk2p. The enzymatic Pfk1-Pfk2 protein complex is shown to the left as a tetramer for simplicity; allosteric adenosine phosphate binding is indicated only (ATP required for catalysis is not shown). The R-state refers to high enzyme activity; the T-state to low activity. Cellular energy levels are indicated with a colour bar (AMP/ADP - ATP levels) to the left. Mg-ATP binding enables Pfk2p-RNA and ribosome interactions enhancing mRNA translation, possibly promoting cell-cycle progression (shown to the right). Parts created with BioRender.com.

### Limitations of the study

Our study concentrated on the yeast *Saccharomyces cerevisiae* Pfk proteins, defining RNA-related functions for the Pfk2p paralog, while such functions for the Pfk1p paralog - if any - remain to be determined. Furthermore, since PFK proteins from diverse species have been shown to bind RNA, it will be of interest to investigate whether the described yeast Pfk2p RNA-binding preferences and RNA unwinding activity is evolutionary conserved and possibly associated with metabolic diseases and/or cancer.

Despite our intense efforts, we have not been able to generate RNA-binding mutants of Pfk2p, which could allow for distinction between RNA binding and enzymatic activity. Hence, a systematic mutational approach based on structural data of Pfk2p in complex with RNA will be of prime interest to deepen our understanding of underlying RNA-protein interactions and to pinpoint relevant domains for RNA unwinding activity; and to solidify the proposed involvement for translation of mRNA targets and cell-cycle progression. Regarding the latter, most of our experiments were performed in asynchronous cell populations, while it is unclear whether RNA-binding activity could depend on a particular phase in the cell cycle. This seems important because oscillations between reductive and oxidative metabolism during mitosis have been observed, opening the possibility that Pfk2p might alternate its roles between mRNA binding and glycolysis as cells transition through various phases of the cell cycle.

## Resource availability

RNA-seq data is available at GEO with access. No GSE151522. All MS files associated with this manuscript are accessible at MassIVE (https://massive.ucsd.edu) under accession number MSV000096586, and deposited at the ProteomeXchange consortium with access. No. PXD058560.

## Acknowledgments

We are grateful to James Cork and Oliver Freeman for help with RNA binding assays, Joe Bryant and Helen King for conducting RT-qPCR measurements, Christina Merk (University of Ulm) for help with the preparative purification of recombinant Pfk proteins, and members of the Gerber lab and Systems Biology tech team for general support and inspiring discussions. We also thank Dr. Martin Pool (University of Manchester) for providing Rpl35 antibodies, and Dr. Matthias Seedorf (University of Heidelberg) for Scp160 antibodies. This study was funded by grants from the Biotechnology and Biological Sciences Research Council (BB/N008820/1, BB/S017747/1) and a Royal Society Wolfson Research Merit Award (WM170036) to A.P.G. T.S. was supported by Cancer Research UK (A29252) and a BBSRC grant (BB/Y004248/1) to M.B.

## Author contributions

Conceptualization, W.S.A, A.M.M.G, and A.P.G; methodology, W.S.A., A.M.M.G, M.B^1^., D.N., T.S. J.J.H., and A.P.G.; formal analysis, W.S.A, A.M.M.G, A.S., and A.P.G; investigation, W.S.A, A.M.M.G, T.S., W.M., A.S., J.H.H., and A.P.G.; resources, M.B^3,5^., D.N., A.S., J.J.H., and A.P.G.; data curation, W.S.A., and A.S.; writing original draft, W.S.A., A.M.M.G, and A.P.G; Visualisation, W.S.A, A.M.M.G, T.S., and A.P.G; project administration, A.P.G.; Funding acquisition, A.P.G.

## Materials and Methods

### Oligonucleotides

DNA and RNA nucleotides are displayed in Table S5.

### *S. cerevisiae* strains and media

Haploid BY4741, BY4742 and diploid BY4743 wild-type (wt) strains, the BY4741 derived *PFK1 (YGR240c)*, *PFK2 (YMR205c)* and *MAP1* (YLR244c) gene deletion strains ^43^, as well as tandem-affinity purification (TAP)-tagged strains ^44^ were obtained from Open Biosystems/ EUROSCARF. Heterozygous *PFK1:TAP*, *PFK2:TAP* diploid strains were generated by mating the BY4741 (*MAT*a) derived *PFK1:TAP* and *PFK2:TAP* haploid cells with wt *MAT*α (BY4742) cells in yeast-peptone-dextrose (YPD; 1% yeast extract, 2% peptone, 2% D-glucose) media ^45^. Heterozygous diploids were selected on synthetic complete media lacking histidine (SC-His) and confirmed by PCR ^46^. Unless otherwise stated, cells were grown in YPD media at 30 °C with constant orbital shaking at 220 r.p.m to mid-log phase (Optical density (OD) at 600 nm ∼ 0.6).

### Plasmids and cloning

CEN/ARS based plasmids for expression of *PFK* wild-type genes (YCplac111:PFK1 (pJJH2101); YCplac111:PFK2 (pJJH2105)) and those with mutations in the Pfk2p catalytic Mg-ATP binding site (YCplac111:*PFK2*^D301T^ (pJJH2504)) and at a proton acceptor for F6P (YCplac111:*PFK2*^D348S^ (pJJH2892)) are based on previously described clones ^19^. The *PFK2* double mutant (DM) at both active sites was constructed by *in vivo* recombination, yielding YCplac111:PFK2^D301TD348S^ (pJJH3063). Therefore, the *EcoR*I site in the polylinker of pJJH2105 was first removed, leading to plasmid pJJH2912. The *EcoR* I digested and dephosphorylated plasmid (pJJH2912) was then co-transformed with a 786 bps DNA fragment (ThermoFisher’s “Gene Art” sting synthesis) covering the two mutation sites and overlapping the two internal *EcoR* I sites. Positive clones were selected on SC-Leu, and plasmids recovered from yeast and transformed into *E. coli* for further propagation and validation by sequencing.

To generate *E. coli* expression constructs, the CDS of *PFK2 and PFK1* were subcloned from the respective pBG1805-PFK1/2 plasmids ^47^ into pDONR221 by Gateway cloning (ThermoFisher, 12536017). The CDSs were then subcloned into the pDEST17 gateway vector (ThermoFisher, 11803012) using LR clonase II kit to generate pDEST17-Pfk1 and pDEST17-Pfk2 expression vectors (ThermoFisher, 12538120). A corrected stop-codon at the end of the CDS of *PFK1* and *PFK2* was inserted using the Q5 site-directed Mutagenesis Kit using primers designed with NEBaseChanger according to manufacturer’s instructions (NEB, E0554S). All constructs were validated by sequencing.

### RNA-binding protein immunoprecipitation (fRIP)

Heterozygous *PFK1:TAP*, *PFK2:TAP*, and wild-type (BY4743; mock control) diploid cells were grown in 400 mL of YPD at 30 °C to mid-log phase (OD_600_∼0.6). RNAs and proteins were crosslinked by the addition of 0.3% formaldehyde (v/v) (ThermoFisher, 28908) to the culture for 10 min at room temperature (RT) upon vigorous shaking of the culture every 2 min. The crosslinking reaction was subsequently quenched by the addition of 125 mM glycine (pH 7.0) for 5 min at RT and constant agitation of cells. Cells were harvested by centrifugation at 2,000 × *g* for 5 min at RT, washed twice with ice-cold phosphate-buffered-saline (PBS) and snap-frozen in liquid nitrogen. The frozen cells were lysed by grinding in a mortar filled with liquid nitrogen and resuspended in 3.5 mL of ice-cold fRIP lysis buffer (20 mM Tris-HCl, pH 7.5, 150 mM NaCl, 500 mM LiCl, 2 mM EDTA, pH 8.0, 5 mM DTT, 0.2% SDS, 1% Triton X-100, 1 × cOmplete™ ULTRA Tablets, Mini EDTA-free (Merck, 05892791001), 1 mM PMSF and 0.2 UmL^-1^ SUPERase•In™ (Invitrogen, AM2696)). Cell lysates were clarified by four subsequent centrifugations at 12,000 × *g* for 10 min at 4 °C. Protein concentrations of extracts were determined with the Pierce BCA assay kit using BSA as a reference standard (ThermoFisher, 23227). Total RNA was isolated by phenol/chloroform extraction and precipitated with ethanol.

The crosslinked TAP-tagged proteins were captured from the extract as follows: 350 µL of Pan mouse IgG Dynabeads (Invitrogen, 11041) were equilibrated in 1 mL fRIP lysis buffer at 4 °C. 2.5 mg of cell extract was then added to the beads and mixed on rotator for 3 h at 4 °C. Beads were collected with a magnet and supernatants kept for monitoring the IP efficiency by Western blot. Beads were thoroughly washed five times for 1 min with 1 mL of high salt buffer (20 mM Tris-HCl (pH 7.5), 500 mM NaCl, 500 mM LiCl, 2 mM EDTA, 0.2% SDS, 1% IGEPAL, 1× cOmplete™ ULTRA Tablets, Mini, EDTA-free, 1 mM PMSF, 0.2 U mL^-1^ SUPERase•In™); and a further five times with 1 mL of 1× AcTEV reaction buffer (50 mM Tris-HCl (pH 8.0), 0.5 mM EDTA (pH 8.0), 5 mM DTT, and 0.2 U mL^-1^ SUPERase•In™). 10 µL (∼3%) of the washed beads were collected for monitoring IP efficiency. To elute Pfk-RNA complexes, the beads were resuspended in 400 µL of 1× AcTEV reaction buffer containing 50 U of AcTEV protease (Invitrogen, 12575-023) and 0.2 U mL^-1^ SUPERase•In™ and incubated for 2 h at RT. The beads were collected with a magnet and the supernatant (= eluates) transferred to a new microtube.

To reverse crosslink and digest proteins, 400 µg Proteinase K (Invitrogen, AM2546) was added to the eluate supplemented with 250 mM NaCl and 0.5% SDS, and incubated for 2 h at 65 °C. RNA was isolated by phenol/chloroform extraction and precipitated with ethanol upon addition of 3 µg of linear polyacrylamide (Invitrogen, AM9520) for at least 1 h at -80 °C. The RNA pellet was washed twice with 70% ethanol, resuspended in 20 µL 1× DNase reaction mix containing 1 µL TURBO DNase (Invitrogen, AM1907)), and incubated at 37 °C for 30 min to digest contaminant DNA. RNA was finally ethanol-precipitated and resuspended in 20 µL of nuclease-free water.

### Reverse transcription (RT)-PCR and sequencing library preparation

For fRIP experiments, 15 μl (75%) of fRIP-RNA and 250 ng of ribosomal RNA (rRNA) depleted total RNA (Ribo-Zero Gold rRNA Removal Kit; Illumina, MRZY1306) was used for reverse-transcription with the SensiFast cDNA synthesis kit containing a mixture of oligo-dT and random hexamers primers (Bioline, BIO-65053). The complementary DNA (cDNA) was further converted to double-stranded cDNA (dscDNA) with the NEBNext® Ultra II Non-Directional RNA Second Strand Synthesis Module as described in the manufacturer’s guide (New England Biolabs, E6111S). DscDNA was sheared to 150 bp fragments by sonication (Covaris S220) with the following settings: Peak incident power: 175 W, Duty Factor: 10%, Cycle per burst: 200 and treatment time: 600 s (10 min). Size distribution of dscDNA was validated with the Bioanalyzer DNA high sensitivity kit (Agilent, 5067-4626). Sequencing libraries were constructed using JetSeq DNA library preparation kit (Bioline, BIO-68025). DNA was sequenced at the Bauer Core Facility (Harvard University, USA) using one lane of HiSeq2000 flow cell with 50 bp single-end reads with a minimum depth of 5 mio reads *per* library.

To validate selected mRNA targets from independent fRIP experiments, RT was performed as described above with 75% of the IPed RNA and corresponding total RNA. 20 µL PCR reactions were performed with 1 μl (5% vol.) of complementary DNA (cDNA), 0.5 µM of each primer (Table S5) and 1 × MyTaq Red Mix (Meridian Bioscience, BIO-25044). The following temperature profile was applied: 5 min at 95°C; 35 cycles of the sequence [95 °C for 30 s, 58 °C for 30 s, 72 °C for 30 s], and 7 min at 72 °C. The products were resolved and visualised on a 2% agarose gel.

### fRIP-seq data analysis

The quality of reads of the RNA-seq data was checked with fastqc v0.12.1 (Babraham institute; https://www.bioinformatics.babraham.ac.uk/projects/fastqc/). Low quality reads were trimmed using FastX toolkit v0.0.14 (Hannon’s lab; http://hannonlab.cshl.edu/fastx_toolkit/) with minimum phred33 score of 20 and minimum length threshold of 20 bp. Total RNA, Pfk1 and Pfk2 fRIP and mock IP reads were mapped to the yeast reference genome R64-2-1 with GMAP/GSNAP ^48^ allowing a maximum of 3 mismatches and output in BAM format. BAM files were sorted and indexed using SAMtools ^49^. Mapping statistics were checked with SAMtools subcommand “flagstat”. Unmapped and low mapping quality (MAPQ) reads were removed from BAM files using SAMtools view. Mapped reads were counted on all genomic features using BEDtools subcommand “multicov” ^50^. Read counts transformed using the variance stabilising transformation method (VST) ^51^. VST-transformed total RNA, fRIP and Mock IP were normalized using trimmed means of M-values (TMM) with limma algorithm ^52^. The Mock control normalised reads were subtracted from fRIP-seq normalised reads of the log_2_ transformed data (log_2_ (fRIP) – log_2_ (mock IP)). *P*-values were calculated using moderated *t*-statistic test and adjusted with Benjamini-Hochberg to obtain false discovery rates (FDR). Significantly enriched RNA targets were at least 4-fold (log_2_ (fRIP) – log_2_ (mock IP) ≥ 2) enriched with ≤ 5% FDR. Processed data is given in Table S1.

### GO analysis and sequence motif searches

Gene ontology (GO) analysis was performed with the Singular Enrichment Analysis (SEA) tool in AgrioGO v2.0 database ^53^ using the *Saccharomyces cerevisiae* dataset. The coding sequences (CDS) of the top 881 Pfk1 and all 867 Pfk2 selected mRNA targets were retrieved from ENSEMBL with BioMart (https://www.ensembl.org/info/data/biomart/index.html). The associated 3’UTR and 5’UTR sequences were extracted from *Saccharomyces* Genome Database (SGD; ^29^ and duplicated and sequences shorter than 10 nts removed, resulting in 875 and 756 3’-UTRs, and 869 and 752 5’-UTR sequences for Pfk1 and Pfk2 mRNA targets, respectively. *De novo* motif searches were performed with MEME version 5.5.1 ^54^ with the following settings: searching the sense strand, RNA sequence, motif width between 6 - 12 nts, minimum and maximum number of sites 20 - 500 per sequence, any occurrence mode and default *p*-value cut-off < 0.0001. To estimate the occurrence of motifs in the background/population; the MEME output motifs were transferred into FIMO implemented in the MEME suite and searched against all CDS (6,255 sequences), and the available 5,211 5’UTR and 3’UTR sequences by searching the given strand only. Motif enrichment was calculated to the respective background/population of sequences, and *p*-values determined (hypergeometric test). MEME search outputs and statistics are given in Table S2.

### RNA-interactome capture (RIC)

Wt (BY4741), *PFK1:TAP*, *PFK2:TAP*, *PGK1:TAP*, *pfk1*Δ and *pfk2*Δ cells were grown in YPD at 30 °C to mid-log phase (OD_600_∼0.6) and collected by centrifugation. Crosslinking, cell collection and lysis were essentially performed as described with fRIP-seq method, except for initial Pfk RIC experiments (Figure 1A) that applied UV-irradiation of cells for crosslinking as described ^55^. Essentially, 300 µL of oligo[dT]_25_ coupled beads (Dynabeads™ mRNA DIRECT™ Purification Kit, Invitrogen, 61011) were equilibrated four times with 1 mL lysis buffer (20 mM Tris-HCl pH 7.5, 150 mM NaCl, 500 mM LiCl, 2 mM EDTA, 5 mM DTT, 0.2% SDS, 1% Triton X100, 1× cOmplete™ ULTRA Tablets, Mini, EDTA-free,1 mM PMSF and 0.2 U mL^-1^ SUPERase•In™). 3 mg of total protein extracts were mixed with beads and incubated at 25 °C for 10 min in a thermomixer (peqlab) at 1,000 r.p.m. For competition experiments, 20 µg of poly(A) was added to extracts (Sigma, P9403) Beads were washed once with 1 mL of wash buffer A (20 mM Tris-HCl pH 7.5, 600 mM LiCl, 0.5 mM EDTA, 0.1% Triton X-100, 1× cOmplete™ ULTRA Tablets, Mini, EDTA-free,1 mM PMSF and 0.2 U mL^-1^ SUPERase•In™), and twice with 1 mL of wash buffer B (20 mM Tris-HCl pH 7.5, 600 mM LiCl, 0.5 mM EDTA, 1× cOmplete™ ULTRA Tablets, Mini, EDTA-free,1 mM PMSF and 0.2 U mL^-1^ SUPERase•In™) each for 15 s. Beads were resuspended in 40 µL of 10 mM Tris-HCl (pH 7.5) and incubated at 85 °C for 5 min. Eluates were collected and stored at -80 °C until use.

### Glucose starvation and polysomal profiling

For standard polysomal profiling, yeast cells were grown in 200 mL of YPD at 30 °C to mid-log phase and treated with 100 µg mL^-1^ of cycloheximide (CHX) for 1 min. Cells were harvested by centrifugation at 1,500 × *g* at RT for 5 min and immediately snap-frozen in liquid nitrogen. For glucose starvation and recovery experiments, 880 mL cultures were grown to mid-log phase. Then 250 mL of cells were removed, CHX treated, collected and snap-frozen in liquid nitrogen. The remaining 550 mL of culture was collected and washed twice with pre-warmed media lacking glucose (YP) and finally resuspended in 550 mL of YP media and grown for 20 min at 30 °C in an orbital shaker. Then, 250 mL of cells were treated with CHX for 1 min and collected. The remaining 300 mL of cells was collected and resuspended in 300 mL of pre-warmed YPD to activate recovery from glucose starvation and grown for 20 min at 30 °C. Finally, cells were treated with CHX, and 250 mL collected and snap-frozen as described above.

Cell pellets were ground in liquid nitrogen using pre-chilled mortar and pestle and resuspended in 2 mL polysome lysis buffer (PLB; 20 mM Tris-HCl, pH 8.0, 140 mM KCl, 5 mM MgCl_2_, 1% Triton X-100, 0.5 mM DTT, 100 µg mL^-1^ cycloheximide, 1× cOmplete™ ULTRA EDTA-free,1 mM PMSF and 0.2 U mL^-1^ SUPERase•In™). For ribosome disassembly experiments, 30 mM EDTA (pH 8.0) was added to extracts in lysis. The cell lysates were centrifuged three times at 16,000 × *g* at 4 °C for 10 min. 5 mg of the resulting extract (total protein) were loaded on top of a 10–50% linear sucrose density gradient prepared in PLB without Triton X-100. Samples were centrifuged at 125,000 × *g* for 2.5 h at 4 °C in a Beckman Coulter Optima XPN-100 ultracentrifuge with SW41Ti swing rotor. Fractions (900 µ) were collected while continuously recording the absorbance at 254nm (A_254_) with a flow cell UV detector (Teledyne ISCO). The ratio (P/S) of polysomal (fraction 7-13) to subpolysomal peak areas (fractions 1-6) was quantified with imageJ.

Total RNA was isolated with Trizol (ThermoFisher, AM9738) and ethanol precipitation. Essentially, 250 µL from each fraction was diluted with 250 µL (1 vol) of cold nuclease-free water, and 10 ng of a control RNA (LysA) was added prior organic extraction. RNA was precipitated by addition of 3 µg of linear polyacrylamide (Invitrogen, AM9520), 0.25 vol. of 10 M ammonium acetate, and 2.5 vol. of ethanol; and RNA pellets washed twice with 70% ethanol, air-dried and resuspended in 20 µL of nuclease-free water. RNA was treated with 1 µL of DNase (TURBO DNA-free, Ambion #1907) prior reverse transcription.

For SYBR Green real-time PCR analysis, RT was performed with 3 µL of RNA from each fraction with the SensiFast cDNA synthesis (Bioline, BIO-65053). qPCR was then performed with 1 µL of the cDNA (accounting for between 30 to 400 ng of RNA extracted from polysome fractions) with the SensiFast SYBR lo-ROX mix (Meridian Bioscience, BIO-94005) according to manufacturer’s instructions. The temperature profile used in qRT-PCR was 95 °C for 2 min, followed by 40 cycles of [95 °C for 5 s, 60 °C for 10 s, 72 °C for 5 s]. Ct values were normalised to the LysA spike-in control as previously described ^56^.

### Immunoblot analysis

0.1% (∼0.7-5 μg of protein) of the input extract, 10% of fRIP eluates, 10% of RIC eluates, and 5 µL of each polysomal fraction were typically resolved by SDS polyacrylamide gel electrophoresis (SDS-PAGE).

Protein samples were resolved on 4–15% Mini-PROTEAN® TGX™ Precast Protein Gels (Bio-Rad, 4561084) and transferred to nitrocellulose (NC) membranes (Cytiva, 10600003) with a semi-dry electrophoretic transfer cell (Trans-Blot Turbo, Bio-Rad). Membranes were blocked in PBS containing 0.1% Tween-20 (PBST) and 5% skimmed milk, probed with designated antibodies and horseradish peroxidase (HRP)-coupled secondary antibodies. Membranes were developed with the Immobilon Western Chemiluminescent HRP Substrate (MerckMillipore, WBKLS0500) and luminescence recorded with a Fusion FX gel documentation system (Vilber Lourmat Sté). The following antibodies were used: peroxidase anti-peroxidase complex (1:5,000; Sigma, P1291) for detection of the TAP-tag, mouse anti-Pab1 (1:5,000; Antibodies online, ABIN1580454), mouse anti-Act1 (1:2,500; MP Biomedicals, 0869100), rabbit anti-PFK1 (1:20,000; ^41^, rabbit anti-Rpl35 (1:5,000, ^57^), rabbit anti-Scp160 (1:10,000, ^57^), HRP-conjugated donkey anti-rabbit IgG (1:5,000; Cytiva, NA9340V), HRP-conjugated sheep anti-mouse IgG (1:5,000; Cytiva, NXA931). Validations of the commercial primary antibodies are provided on the manufacturers’ website.

### Proteome analysis

100 mL of *pfk1*Δ, *pfk2*Δ. *map1*Δ, and isogenic BY4741 wild-type yeast cells were grown in YPD to mid-log phase and collected by centrifugation. Cells were lysed in 50 μl of lysis buffer (100 mM Tris-HCl (pH 8.5), 10 mM Tris(2-carboxyethyl)phosphine) (TCEP), 1% sodium deoxycholate) by strong ultra-sonication (10 cycles, Bioruptor, Diagnode). Protein concentrations were determined by Pierce BCA assay kit using BSA as a reference standard (ThermoFisher, 23227). Sample aliquots containing 50 μg of total proteins were reduced for 10 min at 95 °C and alkylated with 15 mM chloroacetamide for 30 min at 60 °C. Proteins were digested by incubation with sequencing-grade modified trypsin (2% (w/w); Promega, V5113) overnight at 37 °C. Peptides were cleaned up using iST cartridges (PreOmics, Munich, Germany), according to the manufacturer’s instructions. Samples were dried under vacuum and stored at -80 °C.

Sample aliquots comprising 10 μg of peptides were labelled with isobaric tandem mass tags (TMTpro 16-plex, ThermoFisher Scientific) as described previously ^58^. TMT-labelled peptides were fractionated and pooled into 6 fractions by high-pH reversed phase separation using a XBridge Peptide BEH C18 column (3,5 µm, 130 Å, 1 mm x 150 mm, Waters) on an Agilent 1260 Infinity HPLC system as previously described in ^59^. Peptides were subjected to LC–MS/MS analysis using a Q Exactive HF Orbitrap Mass Spectrometer fitted with an EASY-nLC 1000 (both ThermoFisher Scientific) and a custom-made column heater set to 60 °C using the same LC and MS settings as previously described ^60^. The acquired raw data files were searched against a protein database containing sequences of the predicted SwissProt entries of *S. cerevisiae* (www.ebi.ac.uk, release date 2020/04/21), the six calibration mix proteins ^58^ and commonly observed contaminants (in total 12,882 sequences) using the SpectroMine software (Biognosys, version 1.0.20235.13.16424). Standard Pulsar search settings for TMTpro (“TMTpro_Quantification”) were used and resulting identifications and corresponding quantitative values were exported on the PSM level using the “Export Report” function. Acquired reporter ion intensities in the experiments were employed for automated quantification and statistical analysis using an in-house developed SafeQuant R script (v2.3, ^58^). Processed MS data is available in Table S3; associated GO searches are given in Table S4.

### Expression and purification of histidine-tagged fusion proteins in *E. coli*

Expression vectors (pDEST17-PFK1; pDEST17-PFK2) were transformed into ArticExpress (DE3) RIL competent *E. coli* cells (Agilent, 230193). Cells were usually grown in 2YT media (16 g/L tryptone, 10 g/L yeast extract, 5 g/L NaCl) supplemented with 100 µg mL^-1^ ampicillin (Sigma) at 14 °C to mid-log phase (OD_600_ ∼ 0.4). Expression of His-tagged fusion proteins was induced by addition of 1.25 mM or 0.25 mM of isopropyl-β-D-thiogalactopyranoside (IPTG) for 4 or 18 h, respectively. Cells were collected by centrifugation at 5,000 × *g* for 15 mins at 4 °C, washed with ice-cold PBS and snap-frozen in liquid nitrogen and stored at -80 °C.

For small-scale experiments, cells from 100 mL cultures were resuspended in native binding buffer (NB; 50 mM NaH_2_PO_4_, pH 7.5, 500 mM NaCl, 8 mg Lysozyme (ThermoFisher, 89833), 0.01% NP-40, 10 U mL^-1^ DNase I (Promega, M6101), 1 × cOmplete EDTA-free protease inhibitor (Roche, 11873580001)), incubated for 30 min on ice; and cells were lysed with a Sonicator (MSE Sanyo Soniprep 150; 5 × 10 s, 80% power). The cell lysates were then cleared twice by centrifugation at 15,000 × *g* and combined with 500 µL nitrilotriacetic acid-Ni2+ (NTA) agarose resin (ThermoFisher, R901-01) and placed in a chromatography column (Biorad, 7371022). The resin was washed at least 5 times with 2.5 mL of NB buffer, and His_6_-tagged proteins stepwise eluted with increasing concentrations of imidazole (50 - 500 mM). The eluted proteins were dialysed with a G3 Dialysis Cassette (FischerScientific, 10717975) to remove imidazole and changing buffers. Proteins were used for REMSA and initial RNA UV-crosslinking experiments.

To obtain preparative samples, cells from 2 l cultures were resuspended in lysis buffer (LS; 50 mM HEPES, pH 7.5, 1 M NaCl, 10 % Glycerol, 2 mM MgCl_2_, 0.05 % Tween-20, 10 mM Imidazole, 1 mM β-Mercaptoethanol (β-Me)) and disrupted by four to six passages at 1,200 bar through a Microfluidizer (Microfluidics, LM10). Cell lysates were cleared by centrifugation at 30,000 ×*g* for 45 min at 4 °C and loaded on a 5 mL HisTrap FF column (Cytiva, 17531901). The column was washed with 10 column volumes (CV) of high-salt buffer (HS, NB2 buffer with 2 M NaCl, 20 mM imidazole) and 5 CV of LS buffer; and His-tagged proteins were eluted with a linear gradient to 100% elution buffer (EB; 20 mM HEPES, pH 7.5, 0.3 M NaCl, 10 % Glycerol, 2 mM MgCl_2_, 500 mM Imidazole, 1 mM β-Me). Fractions containing recombinant proteins were pooled and dialysed in buffer A (20 mM HEPES, pH 7.5, 150 mM NaCl, 5 % Glycerol, 2 mM MgCl_2_, 1 mM β-Me) and loaded on a HiTrap Heparin HP column (Cytiva, 17040601).

The column was first washed with 10 CV of buffer A. Bound protein was eluted by applying a linear gradient from buffer A to buffer B (containing 2 M NaCl) over 100 mL. Fractions with recombinant protein were pooled and loaded on a HiLoad 16/600 Superdex 200 (Cytiva, 28989335) size-exclusion column (SEC) and separated with buffer A at 1 mL/min. Protein concentrations were determined using a Nanophotometer (Implen, NP80). Required theoretical molecular weights and extinction coefficients were calculated using ProtParam (Expasy). Preparative samples were used for UV-crosslinking and RNA unwinding assays.

### UV-crosslinking assays

Fluorescently labelled RNA oligos with IRDye® 800CW (LI-COR, Inc, USA) were obtained from Integrated DNA technologies (IDT) (Table S5). UV crosslinking was performed by combining 500 fmol of fluorescently labelled RNA and 0.5 µg of His_6_-Pfk2 in 10 µL reaction buffer (20 mM HEPES-KOH pH 7.5, 50 mM NaCl, 2 mM MgCl_2_, 5% glycerol, 1 mM DTT). To test nucleotides, 50 mM of the respective nucleotide (Sigma) was incubated in 50 mM MgCl_2_ for 15 min at RT prior addition of 1 µL to the reaction mix (final conc. 5 mM). Mixed reactions were incubated for 30 min at RT and crosslinked at 254 nm on ice with 400 mJ/cm^2^ in a UV crosslinker (Boekel Scientific, 234100). Samples were resolved on 4-15% gradient SDS-PAGE (Bio-Rad) and scanned with an Odyssey CLx scanner (LI-COR, Inc, USA).

### RNA electrophoretic mobility shift assays (REMSA)

REMSA was essentially performed as described ^61^ with minor modifications. The reaction was set in 20 µL by adding 2 µL of 10 × REMSA buffer (400 mM Tris-HCl (pH 7.5), 300 mM KCl, 10 mM MgCl_2_, 50% Glycerol, 0.1% IGEPAL, 10 mM DTT), 1 µL of 100 nM fluorescently labelled RNA oligo (=5 nM), 10 µL of 1:2 serially diluted recombinant His_6_-Pfk2, 1 µL of 100 mM Mg-ATP (final concentration 5 mM, Sigma, A9187). Reactions were mixed by pipetting and incubated at RT for 30 min. Samples were loaded onto 5% native Tris-acetate (TA) polyacrylamide gel and run at 90 V for 45 min. Gel was scanned with Odyssey CLx scanner (LI-COR, Inc, USA). To determine approximate dissociation constants (K_d_), the intensities of the free and uppermost band representing RNA-protein complexes were quantified with ImageJ (v1.54h, https://imagej.net/ij/). Data was exported to GraphPad Prism and K_d_ estimated at protein concentration where 50% of RNA was bound by the protein.

### RNA unwinding assay

Annealing of the RNA substrates and real-time fluorescence-based RNA unwinding assays were performed as described ^22^. Shortly, 50 nM of annealed RNAs substrates were incubated with 1 µM of the indicated proteins in 10 μl assay buffer (20 mM HEPES-KOH, pH 7.5, 100 mM KCl, 1 mM TCEP, 1% glycerol) in 384-well plates and incubated for 30 mins at RT. Reactions were started by the addition of Mg-ATP, or the indicated nucleotides to a final concentration of 5 mM and fluorescent readings taken in a Spark (Tecan) with excitation at 535 nM and emission at 575 nM for 70 min at RT. Data was analysed according to ^22^. For baseline correction, the buffer control trace (negative control) was subtracted from all other reactions, and the data was normalised to the reference, which is the reporter strand only. The initial linear part of the traces was fitted to yield the initial rate of unwinding and converted to 10^6^ min^-1^. Experiments were performed in triplicates and repeated with at least two different preparative batches of recombinant Pfk1 and Pfk2 proteins.

### Enzymatic activity assays

BY4741 and *pfk2*Δ cells harbouring CEN plasmids expressing wild-type and mutant Pfk2p were grown in 5 mL SC-Leu containing 2% glucose (except BY4741, which was grown in SC). Cells were collected by centrifugation, washed once with 3 mL water and once with 3 mL 50 mM potassium phosphate, pH 7.0 (PP) buffer, and the cell pellet snap frozen. Cells were resuspended in 500 µL of PP buffer and broken mechanically with 500 mg of glass beads (0.3 – 0.5 mm in diameter) upon vigorous shaking for 7 min at 4 °C. 500 µL of cold PP buffer was added and the crude extract was cleared by centrifugation at 14,000 ×*g* for 10 min at 4 °C; and the supernatant used in enzymatic activity assays. PFK activity was measured in a coupled enzyme assay resulting in the oxidation of NADH as described ^19^. Essentially,1 – 5 µL of the extract (concentration 2-5 mg mL^-1^) was combined with 700 µL of the reaction mixture prepared in PP buffer and contained 10 mM MgCl_2_, 5 mM fructose 6-phosphate (F6P), 5 µM fructose 2,6-biphosphate (F2,6BP), 1 mM ATP, 0.2 mM NADH, 0.3 Units (U) aldolase, 0.5 U triose-phosphate isomerase, and 5 U of glycerol phosphate dehydrogenase (Boehringer Mannheim). NADH oxidation was measured by following the decrease in absorbance at 340 nM ^19^.

### Yeast cell growth measurements

Yeast cells were diluted from overnight pre-cultures to a starting OD_600_ ∼ 0.1 in 2 mL of YPD. 0.5 mL of culture (triplicates) were transferred into a 48-well Multidish (ThermoFisher, 150787) placed in a plate reader (CLARIOstar, BMB Labtech), and cells were continuously grown at 800 r.p.m and 30 °C. OD_600_ was recorded every 15 min for 48 hours. Data was exported to GraphPad Prism and doubling time was calculated with the non-linear regression curve fit analysis and selecting the least squares fit method.

### Cell synchronisation and flow cytometry analysis

Yeast precultures were diluted to OD_600_ ∼ 0.1 and grown to mid-log phase in YPD at (30 °C). Cells were diluted again in 25 mL YPD to OD_600_∼0.1 and allowed to grow to early-log phase OD_600_ ∼ 0.2. 1 mL of culture was collected and fixed with 70% ethanol to be used as asynchronous control for the Flow Cytometry (FC) experiment. Cell cycle was blocked by adding 20 µg mL^-1^ of the α-factor (Zymo Research, Y1001) mating pheromone for 180 min at 30 °C upon shaking. To release the α-factor arrest, cells were collected by centrifugation (2,000 ×*g*, 3 min, 30 °C), gently washed with 25 mL of pre-warmed (30 °C) YPD, and finally resuspended in 25 mL of pre-warmed YPD. 1 mL of cells were immediately collected and fixed with 1 mL of 70% ice-cold ethanol, corresponding to start of cell cycle measurements (time = 0). The remaining cells were further incubated on the shaker at 30 °C with 200 r.p.m. Likewise, 1 mL of cells were collected every 10 min and immediately fixed with 70% ice-cold ethanol over a time course of 200 min (20 samples). The collected samples were kept in 70% ethanol at 4 °C for further processing. Fixed cells were centrifuged to remove ethanol, washed twice with 1 mL PBS, resuspended in 1 mL PBS and left in a stand for 1 h at RT for rehydration. Cells were then collected by centrifugation and resuspended in 200 μl of RNase A reaction buffer (2 mM Tris-HCl (pH 7.5), 25 µg of RNase A (ThermoScientific, EN0531)) and incubated at 42 °C for 2 h. Cells were collected by centrifugation (3000 ×*g*, 5 min, RT) and the supernatant discarded. The cell pellets were further washed twice with 1 mL of PBS, resuspended in 200 μl PBS supplemented with 100 µg of proteinase K and incubated at 42 °C for 40 min. Finally, cells were pelleted at 2,000 × *g* for 3 min, washed twice with PBS and resuspended in 1 mL of PBS. DNA was stained with 16 µg mL^-1^ propidium iodide (Alpha Aesar, J66584.AB) for 1 h at RT in the dark. Cells were briefly sonicated and DNA content at each time point was determined on 20,000 cells by flow cytometry analysis with an Attune NxT (Life technologies) cell sorter and the following settings: flow rate of 200 µL s^-1^ and YL-2 detector with long-pass filter and maximum absorbance of 620 nm. Data was saved in FCS 3.0 file format and analysed and visualized using FlowJo (BD Biosciences).

## Supplemental Tables

**Table S1. fRIP RNA-seq data and GO analysis.**

(A) Normalised fRIP-seq and total RNA-seq data. The normalised reads for indicated transcript/genes (rows) across replicate (R1, R2) samples of Pfk1, Pfk2, and mock fRIP as well as total RNA-seq data are displayed in different columns. Log_2_ FC, refers to log_2_ fold-change calculated as the ratio (enrichment) of averaged Pfk1 or Pfk2 normalised reads to the mock samples. Adj.*P*, Benjamini-Hochberg adjusted *p-*value (= FDRs); Target: + indicates selected targets with log_2_ FC > 2 and adj.*P* ≤ 0.05.

(B) GO overrepresentation analysis considering common as well as individual Pfk1 and Pfk2 RNA targets. GO analysis was performed with agriGO Version 2.0. (2023-07-27) considering 7,014 SGD annotated genes as the background gene set; all terms with p ≤ 0.05 are displayed. Columns refer to the following (from left to right). Gene target set: Common (grey), Pfk1 RNA targets (yellow), Pfk2 RNA targets (green); GO category: BP (biological process), MF (molecular function), CC (cellular component); GO Identification number; GO term; Query item, the number of targets assigned to the term; Query total, the total number of annotated RNA targets; Bgitem, the number of annotated genes to respective term; Bgtotal, the total number of annotated genes; p-value; FDR, Benjamini-Hochberg adjusted *p*-value; and SDG ID for target genes.

**Table S2. Motifs identified with MEME among Pfk mRNA targets.**

(A) Motif statistics of MEME and FIMO searches for Pfk1p (top) and Pfk2p (bottom) mRNA targets. Statistics are given for identified motifs identified in the CDS, 5’ and 3’ UTR sequences. *P*-values were determined considering the hypergeometric distribution.

(B) MEME output considering Pfk1p mRNA targets. Consensus motif logos are displayed at the bottom of each alignment.

(C) MEME output considering Pfk2p mRNA targets. A peptide motif search is given to the far-right. Consensus motif logos are displayed at the bottom of each alignment.

**Table S3. MS data for *S. cerevisiae* samples.**

MS data for 3,856 proteins identified with FDR < 1% at the protein identification and quantification level. Columns indicate the following (from left to right): SGD ID; ORF name; Uniprot Accession Number; Gene Symbol. For each sample (*pfk1*Δ, *pfk2*Δ, *map1*Δ, wt) the following information is depicted: PSM, the peptide spectrum match of three biological replicates (Rep1, Rep2, Rep3); log_2_ ratio, calculated as the ratio between average of either *pfk1*Δ, *pfk2*Δ or *map1*Δ PSMs to the wt sample; and q-value (FDR).

**Table S4. GO analysis of proteins with altered abundances in *pfk1*Δ, *pfk2*Δ, and *map1*Δ strains.**

GO overrepresentation analysis for proteins that changed in (A) *pfk1Δ,* (B) *pfk2Δ,* and (C) *map1Δ* cells compared wt cells with ≤10% FDR. GO analysis was performed agriGO Version 2.0. (on 2023-07-27) considering all 3,856 proteins identified by MS as the background gene list. GO terms with Benjamini Hochberg (adj. *p*-value) FDR ≤ 5% are displayed. Columns indicate the following (from left to right): GO category; Set of proteins: proteins with decreased levels (‘downregulated’, red box) *vs.* increased proteins levels (‘upregulated’, green box) compared to wt cells; GO Identification number; GO term; Query item, the number of selected proteins assigned to the GO term; Query total, the total number of selected proteins with annotations; Bg item, the number of proteins in the background (Bb) protein list; Bgitem, the number of annotated proteins within respective term; *p*-value; FDR, Benjamini-Hochberg adj. *p*-value; and SDG IDs for selected proteins.

**Table S5. DNA and RNA oligonucleotides**.

---

**Figure S1.**
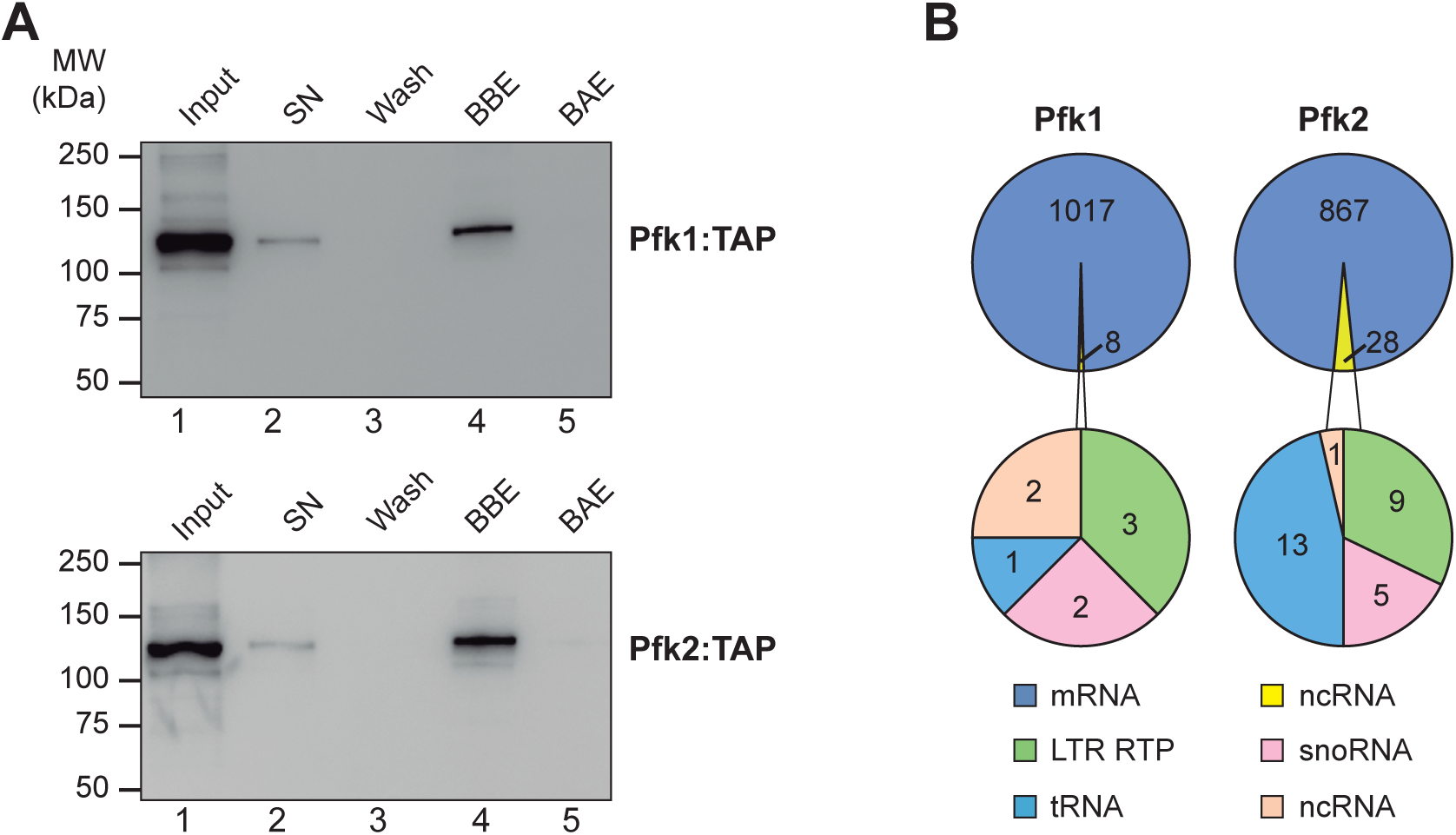
fRIP monitoring and biotype distribution. (A) Immunoblot monitoring the efficiency of Pfk1:TAP and Pfk2:TAP affinity purification. SN, supernatant; BBE, beads before elution; BAE, beads after elution. Molecular weights are indicated to the left. (B) Pie charts showing the distribution of the Pfk1p and Pfk2 fRIP-seq reads on coding and non-coding RNAs.

**Figure S2.**
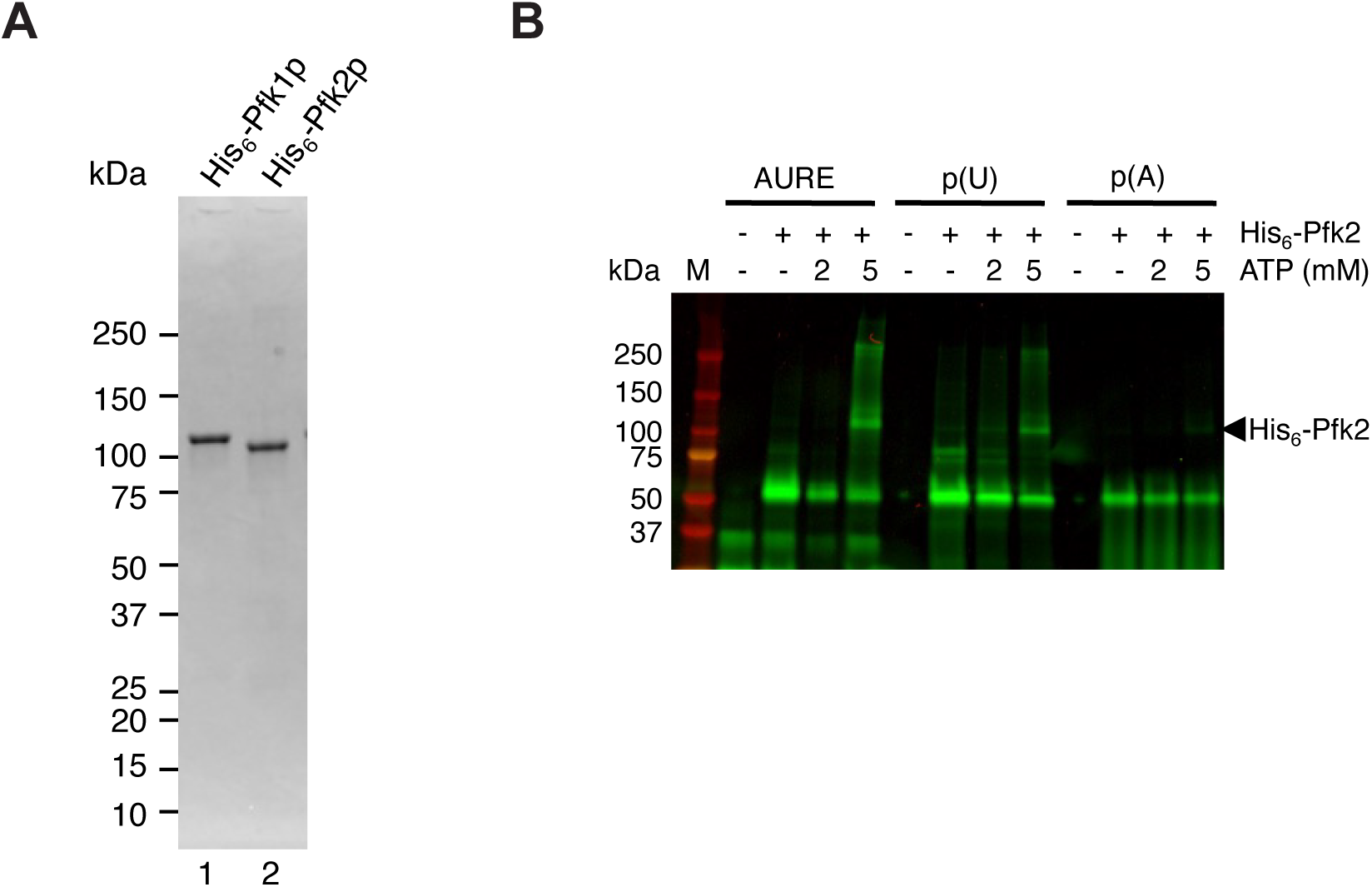
Recombinant Pfk proteins and UV-crosslinking assays. (A) Coomassie staining of recombinant His_6_-Pfk1 (lane 1) and His_6_-Pfk2 (lane 2) proteins expressed and purified from *E. coli*. (B) UV-crosslinking of His_6_-Pfk2 to AURE, poly(U), and p(A) control.

**Figure S3.**
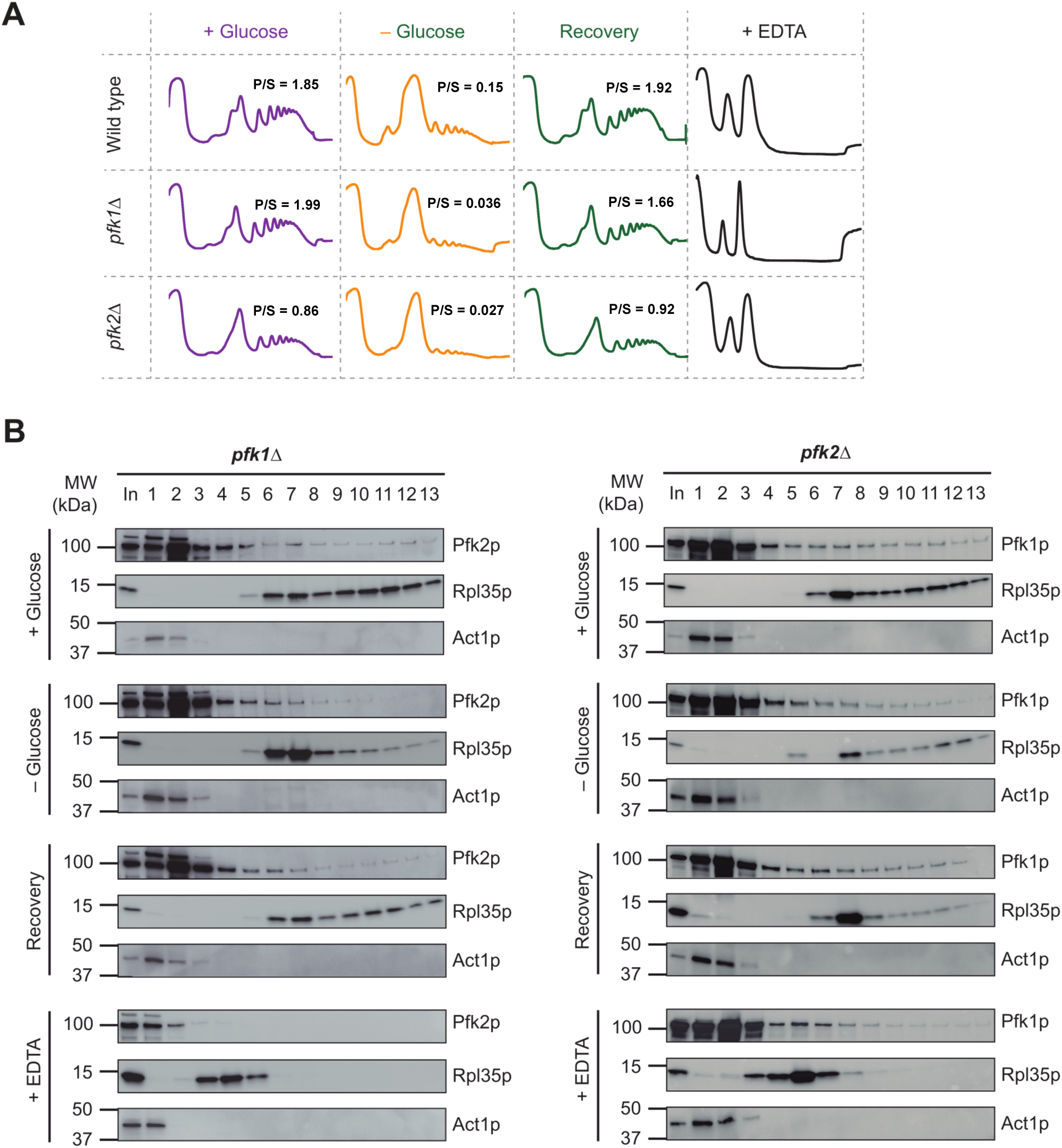
Pfk2p dynamically associates with ribosomes in a glucose-dependent manner. (A) Polysome profiles of wild-type (wt), *pfk1Δ* and *pfk2Δ* strains under normal growth in YPD, glucose (Glc) starvation (-G), recovery conditions (R); and upon disassembly of ribosomes with EDTA (see Figure 3A). P/S indicates the ratio of polysomal (P, fractions 7-13) to sub-polysomal peak areas (S, fractions 1-6 covering free RNPs, 40S, 60S and monosomes). (B) Immunoblot analysis of fraction collected from *pfk1Δ* and *pfk2Δ* mutants. Samples treated with EDTA to dissociate ribosomal subunits was used as control.

